# STING activation reshapes the tumor microenvironment leading to tumor regression in osteosarcoma

**DOI:** 10.1101/2025.10.24.684275

**Authors:** Elizabeth P. Young, Christine A. Johnson, Alex G. Lee, Douglass Morris, Jessica Tsui, Marcus R. Breese, Juan Antonio Camara Serrano, Courtney R. Schott, Clare L. Abram, Stanley G. Leung, Leanne C. Sayles, Aafrin M. Pettiwala, Vanessa S. Gutierrez Vera, Clifford A. Lowell, Alexis J. Combes, Troy A. McEachron, E. Alejandro Sweet-Cordero

**Affiliations:** Division of Pediatric Oncology, University of California San Francisco, San Francisco, CA, USA; Pediatric Oncology Branch, Center for Cancer Research, National Cancer Institute, National Institutes of Health, Bethesda, MD, USA; School of Medicine, University of California San Francisco, San Francisco, CA, USA; UCSF CoLabs, University of California San Francisco, San Francisco, CA, USA; Center for Computational Biology and Bioinformatics, Department of Medical and Molecular Genetics, Indiana University School of Medicine, Indianapolis, IN, USA; Helen Diller Family Comprehensive Cancer Center, University of California San Francisco, San Francisco, CA, USA; Department of Pathobiology, Ontario Veterinary College, University of Guelph, Guelph, ON, Canada; Department of Laboratory Medicine, University of California San Francisco, San Francisco, CA, USA; Division of Radiation Oncology, University of California San Francisco, San Francisco, CA, USA; Department of Pathology, University of California San Francisco, San Francisco, CA, USA; Bakar ImmunoX Initiative, University of California San Francisco, San Francisco, CA, USA; Division of Gastroenterology, University of California San Francisco, San Francisco, CA, USA

## Abstract

Osteosarcomas are characterized by a high degree of aneuploidy, chromothripsis and micronuclei, yet these tumors typically have an immunosuppressive, macrophage-rich, T-cell depleted tumor microenvironment. cGAS-STING dysregulation is a possible mechanism by which immune activation in response to tumor genomic instability could be repressed. We identified almost universal repression of cGAS or STING in human osteosarcomas. However, a STING-activation gene signature was predictive of survival in osteosarcoma patients suggesting potential for activation of this pathway in the osteosarcoma tumor microenvironment. Indeed, in immunocompetent osteosarcoma models, systemic STING agonism led to complete regression and induced lasting immunologic memory. Host STING activation is sufficient to promote this anti-tumor immunity even in the absence of tumor STING. These results nominate the cGAS-STING pathway as an important therapeutic target in osteosarcoma, a disease in which no new curative therapies have been developed in the last 40 years.

**Graphical Abstract:** 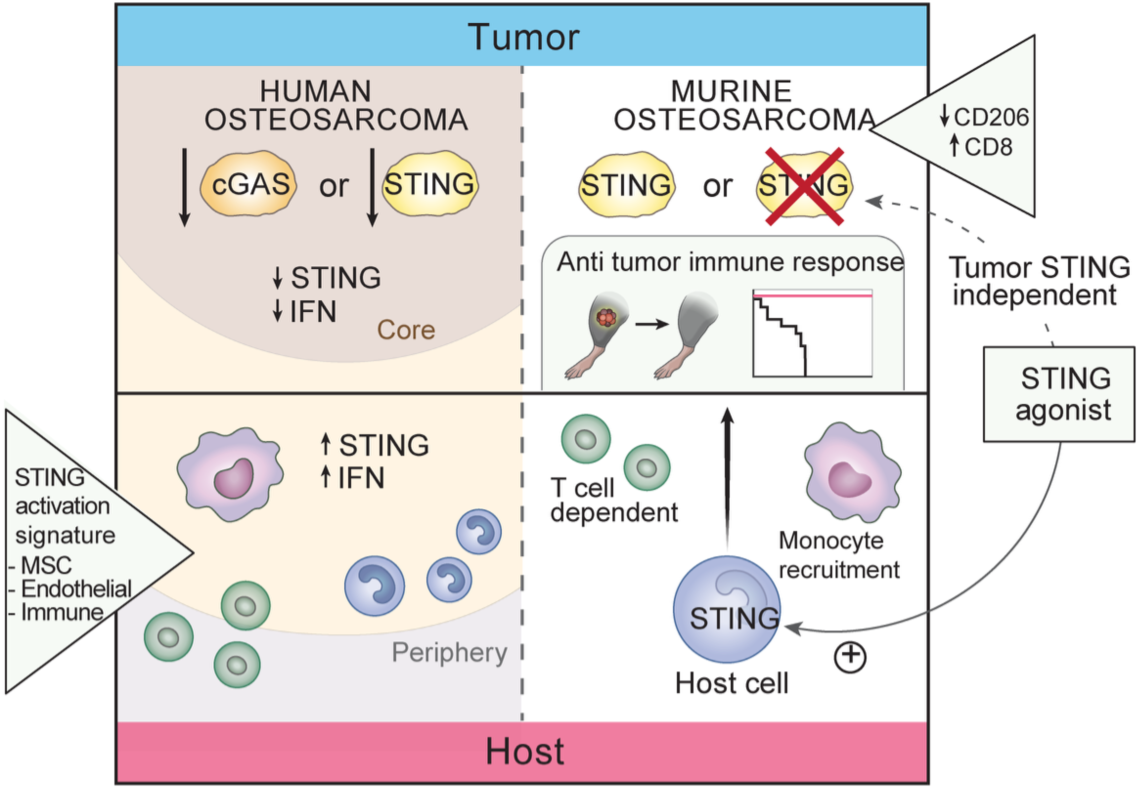

## Introduction

Osteosarcoma is an aggressive malignancy that predominantly affects adolescents and young adults ^1^. Osteosarcoma is characterized by an extremely complex genome with widespread structural rearrangements, aneuploidy and chromothripsis ^2,3^. Despite these massive genomic changes, the osteosarcoma tumor microenvironment is notable for macrophage infiltration and relative T-cell depletion ^4,5^. Identification of effective immune targeting therapies for osteosarcoma and other genomically complex cancers requires an improved understanding of the critical tumor and host factors that drive immune evasion and antitumor immunity ^6–8^. Here we used a combination of human and mouse model systems to characterize the role of cGAS-STING signaling and define the therapeutic potential of activation of this pathway for osteosarcoma treatment.

The cyclic GMP-AMP synthase (cGAS)-stimulator of interferon genes (STING) axis is associated with innate immune activation in multiple cancers ^9,10^. In other genomically complex tumors, chromosomal segregation errors lead to formation of micronuclei resulting in cytosolic DNA and cGAS activation ^11^. Tumor-intrinsic cGAS-STING activation and type 1 interferon production can shape the tumor microenvironment by influencing the recruitment and differentiation of monocytes to pro-inflammatory macrophages ^12^. Repression of cGAS-STING signaling has been demonstrated in several tumor types ^13–15^. The cGAS-STING pathway is attenuated through mechanisms including loss-of-function mutations, hypermethylation of the corresponding promoters, and downregulation of downstream effector signaling ^16,17^. Whether the cGAS-STING pathway is active in osteosarcoma is not well understood.

Given the altered role of the cGAS-STING pathway across many cancers, there is a strong clinical rationale for the development of STING agonists. However, the relative role of tumor-intrinsic vs. non-intrinsic STING activation is not clear and has potentially important implications for development of these agonists. Using a panel of patient-derived (PDX) osteosarcoma cell lines and patient specimens, we evaluated the functional status of the cGAS-STING pathway in human osteosarcoma (**Graphical Abstract**). In almost all cases, there was repression of either cGAS or STING, suggesting a strong pressure to silence this pathway during tumor evolution. RNAseq of osteosarcoma PDX cell lines treated with STING agonist defined an osteosarcoma-specific STING activation signature. Evaluation of primary patient samples for expression of this signature demonstrated a protective effect on progression-free and overall survival. In syngeneic immunocompetent mouse models of osteosarcoma, pharmacologic STING activation had tumor-eradicating effects that did not depend on tumor STING function, nominating a potential broad translational role for this therapy in osteosarcoma patients despite intratumoral cGAS-STING repression. STING agonist treatment establishes long-term immune memory in osteosarcoma, allowing recipient mice to reject tumor rechallenge through a mechanism that is dependent on T cell memory.

## Results

### Integrity of the cGAS-STING pathway in patient-derived osteosarcoma cell lines

We implemented a cross-species strategy to functionally investigate the role of cGAS-STING in osteosarcoma (**Fig. 1A**). Across a panel of human osteosarcoma cell lines, cGAS and STING protein levels were highly heterogeneous (**Fig. 1B**) ^18^. In almost all cases, cell lines with high cGAS had low STING and the inverse was also true (**Fig. 1B**). As *cGAS* and *STING1* mRNA transcripts were detectable in all cell lines this result indicates a post-transcriptional mechanism for repression of these proteins (**Fig. S1A-B**). To determine the capacity for STING pathway activation in these cell lines, we used the non-cyclic di-nucleotide STING agonist diABZI ^19^. Treatment with diABZI induced markers of STING activation, including phospho-STING (pSTING), phospho-TBK1 (pTBK1), phospho-IRF3 (pIRF3), and phospho-STAT1 (pSTAT1) in the STING-high, but not the STING-low cell lines (**Fig. 1C**). STING-high lines also demonstrated a robust induction of *ISG15* and *CXCL10* gene expression as well as protein expression of CCL5 and CXCL10 (**Fig. 1D-E**).

**Figure 1.**
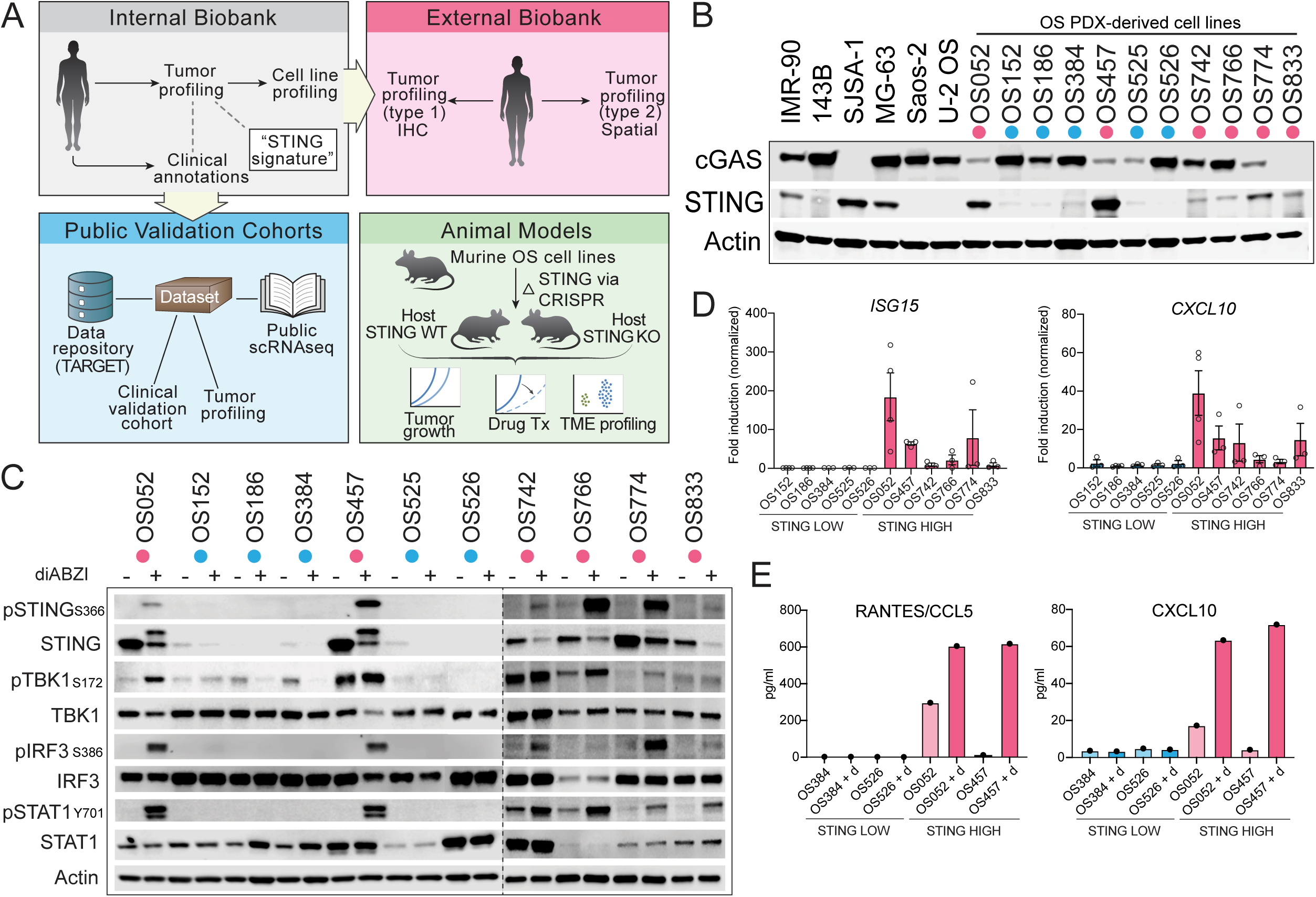
A subset of osteosarcoma patient-derived models demonstrates STING protein repression. **1A.** Schema of models and datasets used. **1B.** Immunoblot for cGAS and STING in osteosarcoma cell lines under standard culture conditions. **1C.** STING effector activity in osteosarcoma PDX cell lines 6h after STING agonist (diABZI). **1D.** RT-qPCR of interferon stimulated genes in osteosarcoma PDX cell lines 24h after diABZI stimulation (mean ± SEM; n=3-4 independent experiments). **1E.** Pro-inflammatory cytokine levels (pg/mL) in conditioned media of STING-high (OS052, OS457) vs. -low (OS384, OS526) osteosarcoma PDX cell lines upon stimulation with diABZI (“+ d”) for 6h (mean; n=2). Cell lines with high vs. low STING protein in 1B are color coded accordingly as STING-high (pink) or STING-low (blue).

DNA-bound cGAS produces cGAMP, which directly activates STING ^20^. To determine whether total cGAS protein correlates with micronuclei (MN) burden and cGAMP production, we quantified cGAS+ MN (MN that co-stain positive for cGAS suggesting co-localization of cGAS to MN) by immunofluorescence and cGAMP production by ELISA. Cell lines with higher cGAS protein showed a trend towards a greater number of cGAS+ MN per total nuclei (**Fig. S1C-D**) as well as higher intracellular cGAMP (**Fig. S1E**-**F**). These findings suggest the existence of two distinct groups of osteosarcoma cell lines: those that are “cGAS-high/STING-low” and those that are “cGAS-low/STING-high.” This in turn suggests a conserved capacity for inflammatory signaling through STING in response to diABZI only in the cGAS-low/STING-high group (**Fig. S1G**).

### Chronic activation of cGAS suppresses STING expression in patient-derived osteosarcoma cell lines

While we did not observe a correlation between *STING1* mRNA transcript and STING protein levels, there was an inverse relationship between STING and cGAS protein levels in these cell lines (**Fig. S1A-B**). As STING protein is known to be actively degraded after cGAMP-mediated activation ^21,22^, we hypothesized that chronic cGAS activation could be responsible for low STING protein levels in this subset of cell lines (**Figure S1G**). To investigate the relationship between cGAS activity and STING repression, cGAS-high/STING-low cell lines were treated with a cGAS inhibitor (G150). Treatment with this inhibitor did indeed lead to an increase in STING protein levels (**Fig. S1H**). However, stimulation of the downstream pathway with diABZI after rescuing STING levels with cGAS inhibition did not result in downstream effector activation (**Fig. S1I**). Similarly, the endo/lysosomal proton pump inhibitor bafilomycin resulted in increased total STING protein levels but did not restore STING activation capacity (**Fig. S1J-K**) ^21^. Together, these data suggest that increased levels of total STING protein are by themselves not sufficient to reactivate STING in osteosarcoma cell lines.

### STING activation signatures indicate improved overall and disease-free survival in patients with osteosarcoma

We defined the effect of diABZI treatment on osteosarcoma gene expression using four STING-high osteosarcoma PDX cell lines (**Fig. 2A**). RNAseq identified a “STING activation” gene signature comprised of 94 consistently upregulated genes (**Fig. 2B; Supplemental Table 1**). As expected, this signature was enriched for pathways involving interferon signaling and inflammatory responses (**Fig. S2A-B**). Given the above observed suppression of cGAS-STING in osteosarcoma cell lines, it was unclear whether this gene signature would be present in human osteosarcoma tumors. To investigate this further, single-sample gene set enrichment analysis (ssGSEA) using this signature was performed on expression data from a cohort of primary osteosarcoma biopsies (**Fig. 2A**; **Fig. 2C-D**). Surprisingly, high expression of the STING activation signature was associated with improved overall and disease-free survival in a cohort of primary osteosarcoma patient samples (n=23). The high STING signature group showed prolonged overall survival (p=0.022, HR=0.138, 95% CI: 0.017-1.128) and disease-free survival (p=0.015, HR=0.18, 95% CI: 0.038-0.858), with an 86.2% and 82% reduction in the risk of death and disease progression, respectively (**Fig. 2C-D**). These findings were corroborated in an independent osteosarcoma dataset (see methods) (overall survival; n=72, p=6.23e-05; HR=0.128, 95% CI: 0.038-0.437; **Fig. 2E**).

**Figure 2.**
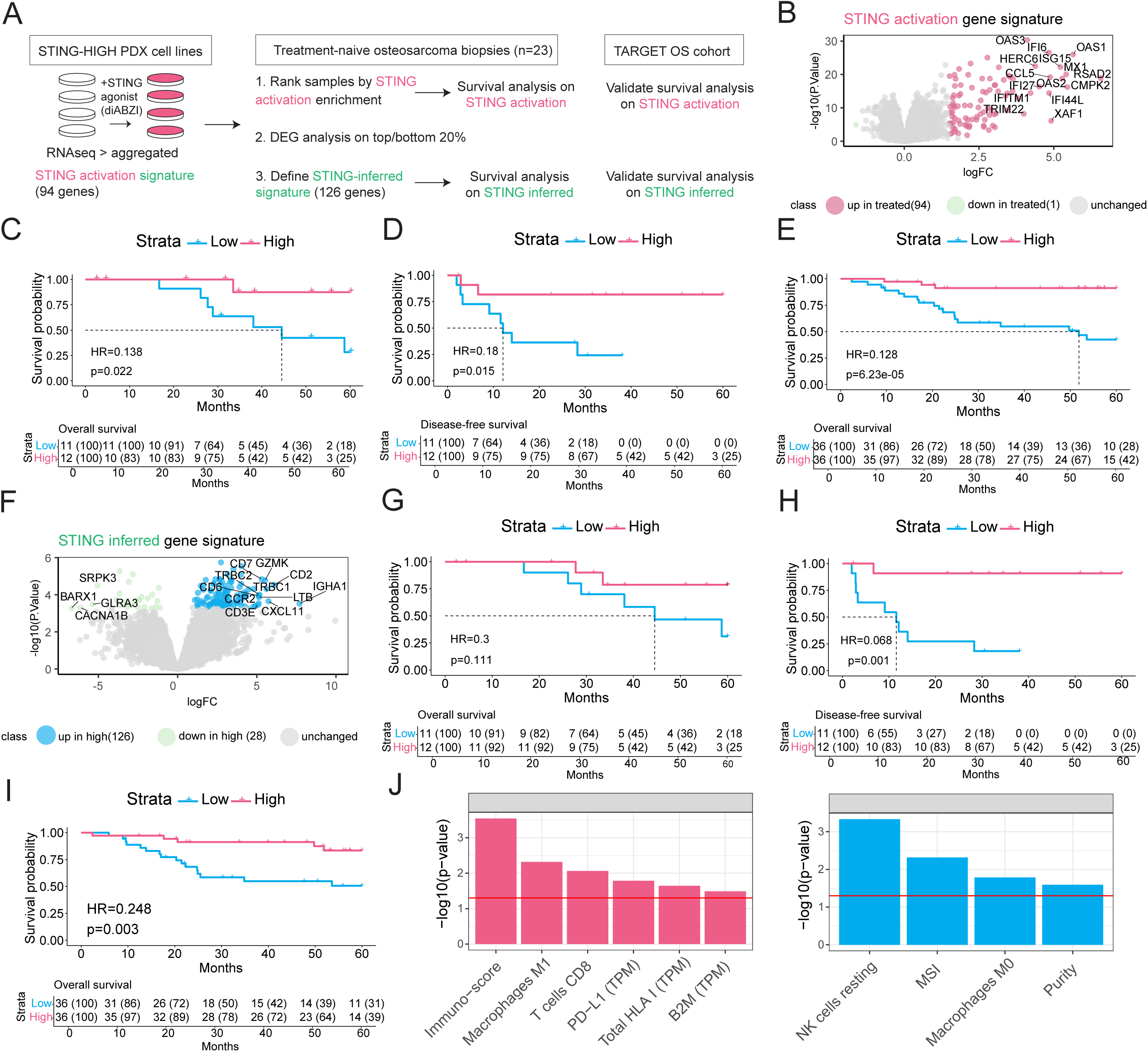
A STING activation gene signature is protective in patients with osteosarcoma. **2A.** Schema. **2B.** STING-high PDX-derived osteosarcoma cell lines (OS052, OS457, OS742, and OS774) were treated with STING agonist diABZI and probed with bulk RNA sequencing to define an aggregate geneset of the transcriptomic changes associated with STING activation in osteosarcoma, termed “STING activation”. **2C-E.** Survival analysis after stratifying samples (n=23) by STING activation signature enrichment (top vs. bottom 50%), with overall (2C) and disease-free survival (2D) and overall survival validation using an independent osteosarcoma dataset (n=72; 2E). **2F.** Biopsy samples (n=23) were stratified by high and low for STING activation signature (top and bottom 20%) and DEG was utilized to evaluate for transcriptomic differences between these cohorts, defining a second signature termed “STING inferred;” genes in original STING activation signature were excluded. **2G-I.** Survival analysis on STING inferred gene signature in 23 diagnostic biopsy specimens, with overall (2G) and disease-free survival (2H) and overall survival validation using an independent osteosarcoma dataset (n=72; 2I). **2J.** Using a generalized linear model, genomic and immune cell features were analyzed to determine how predictive each feature was to the “STING activation” gene expression signature score (z-scaled) (left plot=positive association; right plot=negative assocication). Significance scores were calculated for each feature. Multiple testing correction was performed using Benjamini–Hochberg. Features that had a predictive significance (corrected) less than 0.05 are shown. The red line indicates p=0.05.

Next, we sought to identify additional genes whose expression in tumor samples correlated with that of the STING activation signature (**Fig. 2A**). We reasoned that this strategy would identify genes upregulated in the tumor microenvironment of osteosarcoma tumors in response to STING activation. We identified 126 highly expressed genes we labelled as “STING inferred” (**Fig. 2F; Supplemental Table 2**). Many of the genes in this “STING inferred” signature are associated with innate and adaptive immunity, including cytokines that mediate monocyte recruitment as well as genes responsible for T and B cell function. Ranking diagnostic biopsy samples for enrichment of this signature and stratifying top vs. bottom 50% in an additional survival analysis revealed an association between the STING inferred signature and patient outcomes, with a trend towards improved overall survival (p=0.111; HR=0.3, 95% CI: 0.06-1.487; **Fig. 2G**) and improved disease-free survival (p=0.001; HR=0.068, 95% CI: 0.008-0.538; **Fig. 2H**). This association was also seen in an external dataset (n=72, p=0.003; HR=0.248, 95% CI: 0.091-0.68; **Fig. 2I**).

Next, we assessed the bulk RNA sequencing data to identify correlations with other features of the tumor samples (**Fig. S2C**). Several genomic or tumor microenvironment features were either positively (**Fig. 2J, left**) or negatively correlated (**Fig. 2J, right**) with the STING activation signature, including tumor purity, which likely reflects immune or stromal cell infiltration into tumors. We then used a deconvolution algorithm (CIBERSORTx; see methods) to identify correlation between abundance of specific cell types with high enrichment of the STING activation signature. M1 macrophages, activated NK cells, and CD8+ T cells were all associated with high STING activation signature. Other studies have also found that M1 and M2 macrophages were most abundant in a favorable prognosis group of human and canine osteosarcoma patients, and higher M0 macrophages were observed in a poor prognosis group ^23^.

### Single-cell RNA sequencing defines cell type representation of STING signatures in patient samples

Next, we mined publicly available scRNAseq datasets from primary and metastatic osteosarcoma specimens to examine the enrichment of the STING signatures in individual cell types ^24,25^. These published datasets were re-analyzed and annotated using pipelines for cell marker expression (**Fig. 3A; 3D; Supplemental Table 3**) and copy number inference to distinguish tumor from non-tumor (see methods; **Fig 3B; 3E**). As the “STING activation” signature was obtained from osteosarcoma cells, we hypothesized that it would be most upregulated in tumor cells. Unexpectedly, we observed that this signature was enriched only in a subset of tumors cells. Surprisingly, it was also expressed in multiple other cell types across both datasets (**Fig. 3C; 3F**). In treatment-naïve samples the STING activation signature was enriched primarily in myeloid and tumor-infiltrating lymphocytes whereas in pre-treated samples there was more evidence of STING activation in endothelial cells and osteoblasts (**Fig. 3C; 3F**). In contrast, the STING inferred signature was enriched primarily in myeloid and TIL/NK populations in both datasets, although there was also enrichment in osteoclasts in the treatment-naïve samples (**Fig. 3C; 3F**). Taken together these results support a complex role for STING activation in the OS tumor microenvironment and our observation above that STING is largely silenced in the tumor compartment.

**Figure 3.**
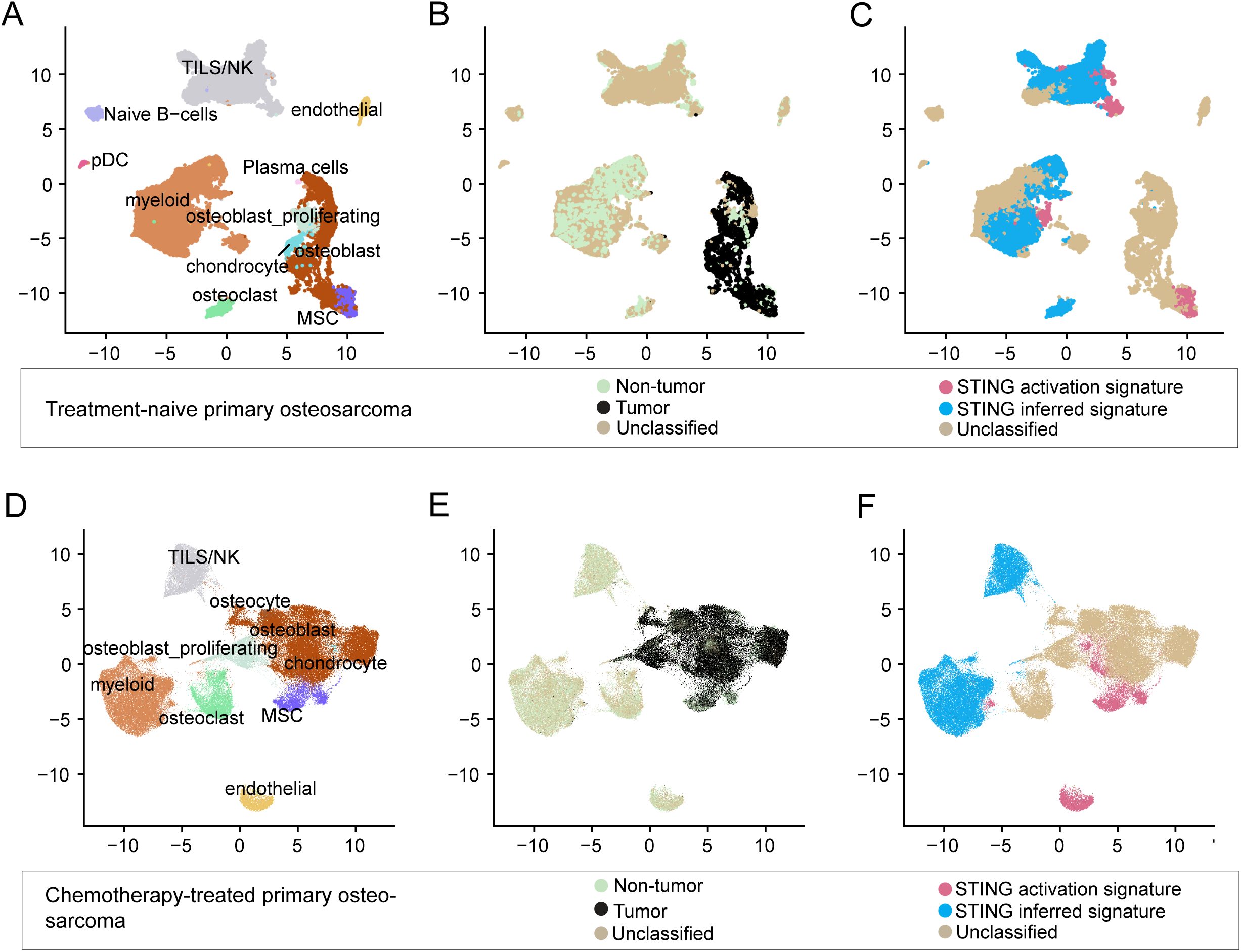
Re-analysis of scRNAseq from 6 treatment-naïve osteosarcoma samples and 11 treated osteosarcoma samples demonstrates cell type contribution to STING signatures. A cohort of 6 treatment-naïve primary osteosarcoma samples is included in **3A-C** (25) and a cohort of 11 treated osteosarcoma samples (9 primary and 2 metastases) is included in **3D-F** (24). **3A.** All cells in all samples annotated by cell-type, see also **3D**. **3B**. All cells in all samples annotated by tumor vs. normal employing inferCNV to infer cells with altered copy number, see also **3E. 3C.** All cells in all samples annotated by enrichment for either STING activation signature as defined by RNAseq experiment performed in Fig. 2B or STING inferred signature as described in text, see also **3F.**

Each dataset was then analyzed on a per-sample basis (**Fig. S3A; S3D** and **Fig. S3B-C; S3E-F**). In the treatment-naïve samples, only a small fraction of tumor cells was enriched for the STING activation signature. In contrast, the STING-inferred signature was more widely expressed with a majority cells expressing this signature in the nontumor fraction (**Fig. S3C**). In the pre-treated samples, STING activation enrichment was variably identified in < 25% of total cells across all samples (**Fig. S3D**). The STING inferred signature was clearly localized to myeloid and lymphocyte populations, confirming that the heuristic for obtaining this signature enriched for genes associated with STING in the tumor microenvironment (**Fig. S3F**).

### Spatially distinct expression of STING and type I interferon signaling in metastatic and non-metastatic osteosarcoma specimens

To evaluate the spatial distribution of STING activation we performed IHC to detect STING protein expression in a cohort of 20 matched primary and metastatic osteosarcoma pairs. Metastatic tumors were split into periphery (non-malignant stromal tissue adjacent to the tumor) or tumor core (region with high tumor cell content). All primary tumor specimens had high tumor cell content and were devoid of adjacent non-malignant stromal tissue. Quantification of STING protein expression revealed similar labeling intensity in the tumor core of primary and matched metastatic tumors (**Fig. 4A-D**). However, STING staining intensity was lower in the core of both primary and metastatic tumors compared to the peripheral regions of metastatic specimens. These data confirm that STING is primarily expressed in immune and/or stromal cells and not within the tumor cells, consistent with the scRNAseq data and the cell line analysis above.

**Figure 4.**
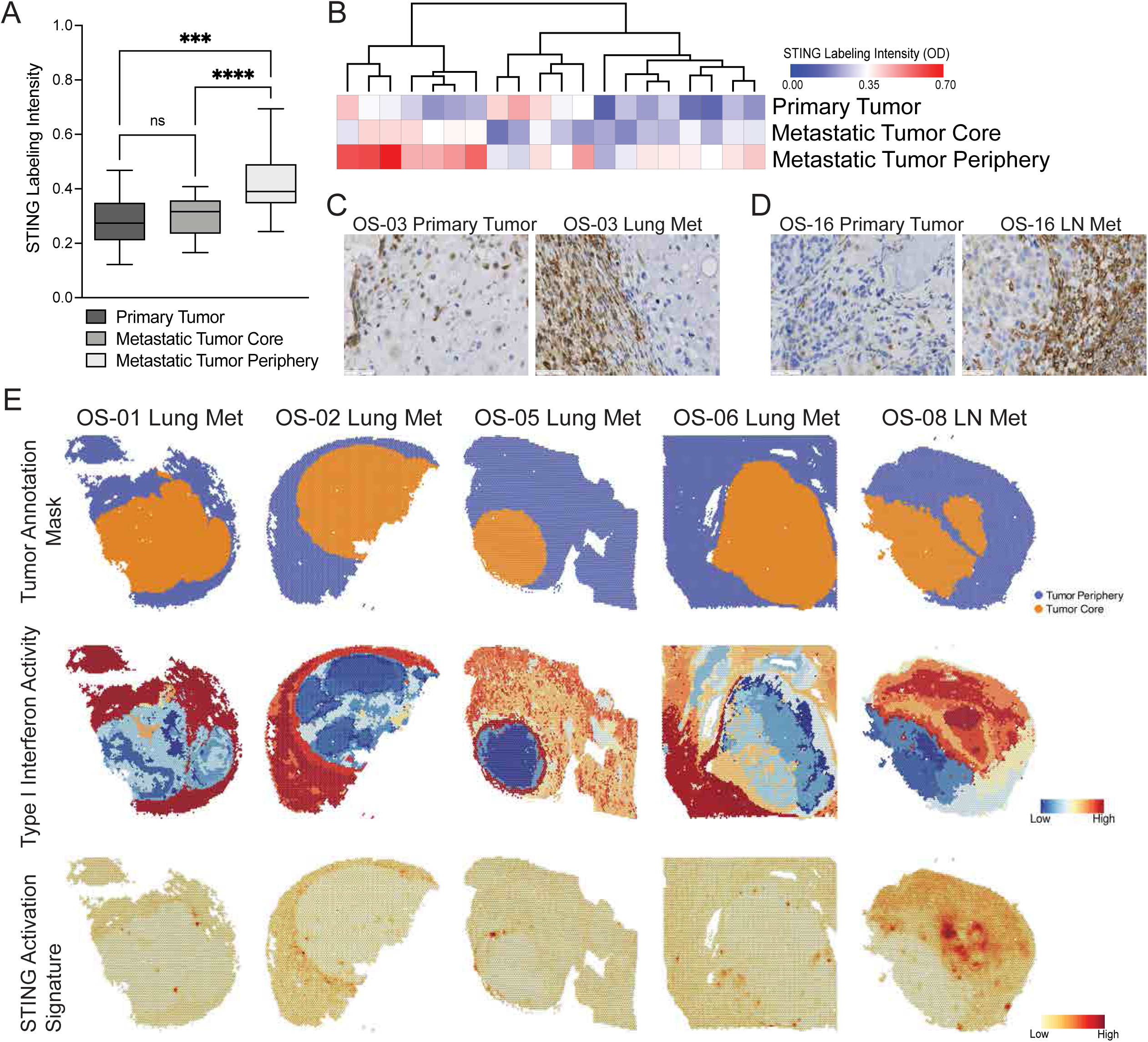
Spatial expression patterns of STING in primary and metastatic osteosarcoma patient specimens. **4A.** Boxplot showing STING labeling intensity (avg. OD) in n=20 matched primary-metastatic specimens. Statistical significance (p<0.05) determined by repeated measures one-way ANOVA. **4B.** Unsupervised hierarchical clustering of STING labeling intensity. **4C-D.** Representative images of STING labeling intensity in (C) the primary tumor core (left) and pulmonary metastatic lesion (right) of specimen OS-03 and (D) the primary tumor (left) and lymph node metastatic lesion (right) of osteosarcoma specimen OS-16. Scale bar = 50 [)m. 4E. Spatial transcriptomic profiling showing regional distribution of type I interferon activity and STING activation signature in metastatic osteosarcoma specimens. Top row: masked spatial plots differentially highlighting tumor core and tumor periphery. Middle row: spatial plots of CytoSig analysis showing the regional distribution and relative strength of type 1 interferon signaling activity. Bottom row: spatial enrichment of STING activation signature.

Next, we evaluated the regional distribution of computationally inferred type 1 interferon activity in metastatic osteosarcoma specimens using spatial transcriptomic profiling (see methods). The type 1 interferon activity signature reflects transcriptional surrogates of downstream signaling activity generated using a combination of experimentally derived signatures and data mining ^26^. Type I interferon activity was predominantly localized to the tumor periphery (**Fig. 4E**). We also mapped the STING activation signature using the gene set described above and identified a similar localization to the tumor periphery (**Fig. 4E**). Taken together, these results suggest that non-malignant cells in the tumor microenvironment are the primary cellular source of STING activity in human osteosarcoma.

### Tumor STING loss has no effect on tumor growth or metastasis

To investigate the effect of STING activation *in vivo*, we used the syngeneic osteosarcoma models K7M2.1 (BALB/c) and F420 (C56BL/6) ^27,28^. (**Fig. S4A**; see methods). Both cell lines have functional STING activation as demonstrated by treatment with either diABZI or the cGAS agonist G3-YSD (**Fig. 5A**). To study the effect of cell-autonomous STING in these models, isogenic STING knock-out cell lines were generated (**Fig. 5B**).

**Figure 5.**
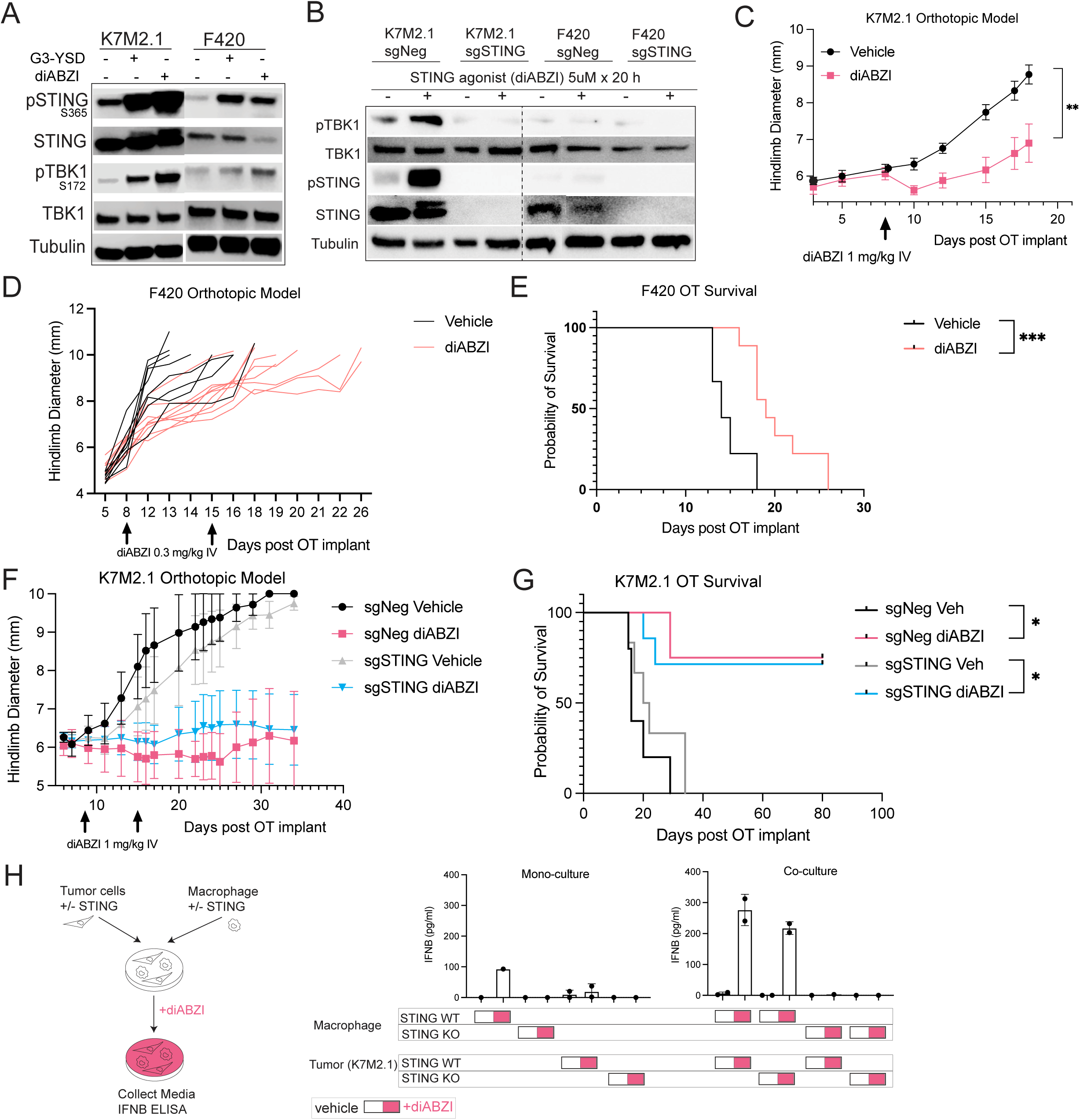
Short-term treatment with systemic STING agonism causes long-term tumor eradication. **5A.** STING activation (pSTING) and downstream STING effector activity (pTBK1) evaluated in murine osteosarcoma cell lines (K7M2.1, F420) 6h after cGAS agonist (G3-YSD) or STING agonist (diABZI). **5B.** pSTING and pTBK1 were evaluated in murine osteosarcoma STING KO cell lines (K7M2.1, F420) 20h after a 6h treatment course with diABZI. **5C.** Tumor growth of K7M2.1 OT implants in BALB/c mice treated with diABZI (1mg/kg IV). Vehicle, n=7; diABZI, n=6. **5D-E.** Tumor growth of individual mice (D) and survival curve (E) of F420 OT implants in B6 mice treated with diABZI (0.3mg/kg IV, weekly x 2). Vehicle, n=8; diABZI, n=9. **5F-G.** Tumor growth (F) and survival curve (G) of K7M2.1 sgNeg and sgSTING OT implants in BALB/c mice treated with diABZI (1mg/kg IV, weekly x 2). sgNeg vehicle, n=5; sgNeg diABZI, n=4; sgSTING vehicle, n=6; sgSTING diABZI, n=7. **5H.** Schema (left) of co-culture. Analysis of IFN-[l production (pg/ml) (right) in conditioned media from BMDMs (STING WT and STING KO) and K7M2.1 isogenic cell line pair (K7M2.1 sgNeg and sgSTING) alone and in co-culture stimulated with diABZI for 6h, assessed at 24h by ELISA (n=2). Data are shown as mean ± SEM (C, F) and mean ± SD (H); *, P<0.05; **, P<0.01; ***, P<0.001 unpaired t test of tumor size at end of study (C), log-rank (Mantel-Cox) test (E, G).

For *in vivo* studies we used an orthotopic (OT) model in which tumor cells are implanted in the hind limb. After tumor growth, the hind limb is amputated and mice are evaluated for spontaneous metastasis to the lung ^18^. An experimental metastasis model using tail vein inoculation of tumor cells was also used for comparison. Growth of K7M2.1 OT primary tumors was not impacted by loss of STING (**Fig. S4B-C**). Furthermore, we did not observe differences in survival between control and STING KO K7M2.1 tumors after amputation of the primary tumor and surveillance for lung metastasis (**Fig. S4D**). This result is in contrast with prior reports in which tumor-intrinsic STING was suggested to play a role in breast cancer metastasis ^29^. This discrepancy may reflect intrinsic differences between tumors of epithelial and mesenchymal origin or perhaps a distinct role for STING in osteosarcoma.

To evaluate the impact of tumor-derived STING signaling on immune cells, OT tumors were isolated at the primary tumor endpoint for flow cytometry immunophenotyping. We did not observe differences in the myeloid (**Fig. S4E, left panel**) or lymphoid (**Fig. S4E, right panel**) compartments (see **Fig. S5A-B** for gating strategy). T lymphoid immunophenotyping revealed a modest decrease in CD8+ T cell abundance in the STING KO tumors, with an altered T cell phenotype (Ly108-CD69+; “Tex term” state), but this did not reach statistical significance. The “Tex term” state refers to terminally exhausted resident T cells suggestive of prior activation ^30^.

We then evaluated the effects of tumor STING KO in an experimental (tail vein) metastasis model. There was no difference in tumor burden or tumor number after inoculation between control and STING KO K7M2.1 tumors (**Fig. S4F**). To ensure the absence of a phenotype with STING KO was not model dependent, these experiments were repeated using the F420 cell line in B6 mice. Again, we did not observe differences between animals bearing control vs. STING KO tumors in time to amputation, survival with a lung metastasis burden endpoint, or tumor microenvironment constitution (**Fig. S4G-J**).

### Tumor STING is dispensable in long-term disease control that results from limited dosing of systemic STING agonist

We then evaluated the effect of the systemically administered STING agonist diABZI in the immunocompetent models ^19^. Animals were randomized to vehicle or diABZI 8 days after K7M2.1 OT implantation. We observed a reduction in OT tumor growth after a single dose of diABZI (**Fig. 5C**). Similarly, two doses of diABZI in the B6/F420 orthotopic tumor model prolonged survival (**Fig. 5D-E**). Next, we used isogenic K7M2.1 STING KO cells in the orthotopic model to evaluate the dependence of the diABZI effect on tumor STING (**Fig. 5F-G**). Control (non-targeting sgRNA) or tumor STING KO cell lines were implanted OT, and mice were randomized to receive vehicle or diABZI 8 days after OT implantation. Two doses of weekly diABZI were sufficient to induce complete tumor regression in 75% of mice regardless of the presence of tumor STING (**Fig. 5F-G**). In mice with tumors that regressed, we observed no tumor regrowth for at least 2.5 months after treatment. Thus, a limited course of diABZI provides a survival benefit that is independent of tumor STING status and in most mice appears to induce a long-lasting complete response.

To further evaluate the role of tumor vs. host STING in the response to STING agonism, we used an *in vitro* co-culture system of tumor cells and macrophages. The K7M2.1 control or STING KO isogenic cell line were co-cultured with bone marrow-derived macrophages (BMDMs) obtained from STING wild-type mice (C57B6) or STING KO mice (C57B6) (**Fig. 5H**). At baseline, IFN-β was undetectable in conditioned media from both tumor cells and BMDMs cultured alone or in co-culture. STING WT BMDMs cultured with tumor cells and treated with diABZI displayed increased IFN-β relative to BMDMs cultured in isolation, independent of tumor STING status, supporting the hypothesis that macrophages respond directly to diABZI via intrinsic activation of STING rather than by activation of tumor STING. However, the higher IFN-β levels in both co-culture settings (with/without tumor STING), compared to either tumor or macrophage monoculture, indicate that interactions between tumor and macrophage facilitate an amplified response to diABZI.

### Systemic STING agonism induces early T cell recruitment followed by myeloid infiltration

We performed time-course experiments to define the immune response to diABZI in K7M2.1 tumor-bearing mice. Mice were randomized to untreated and treatment groups after OT implantation and immunophenotyping was performed at early (2 hours) and late (48 hours) timepoints after a single dose of diABZI (**Fig. 6A-B**). At the early time point, we observed an increase in T cell infiltration in tumors both as a fraction of CD45 and as an absolute count normalizing for weight (**Fig. 6B**). The increase in absolute T cell count per tumor weight persisted after 48 hours, whereas the relative T cell fraction decreased, likely due to infiltration of monocytes and neutrophils at this later timepoint (**Fig. 6B**). Monocytes and neutrophils were not increased either in relative or absolute terms at the early timepoint, suggesting the initial immune response to diABZI results in rapid T cell recruitment that is not initially accompanied by changes in the myeloid fraction. At 48 hours after STING agonism, we also observed a relative decrease in mature myeloid cells with a shift away from TAMs and a trend towards decreased DCs (**Fig. 6B**).

**Figure 6.**
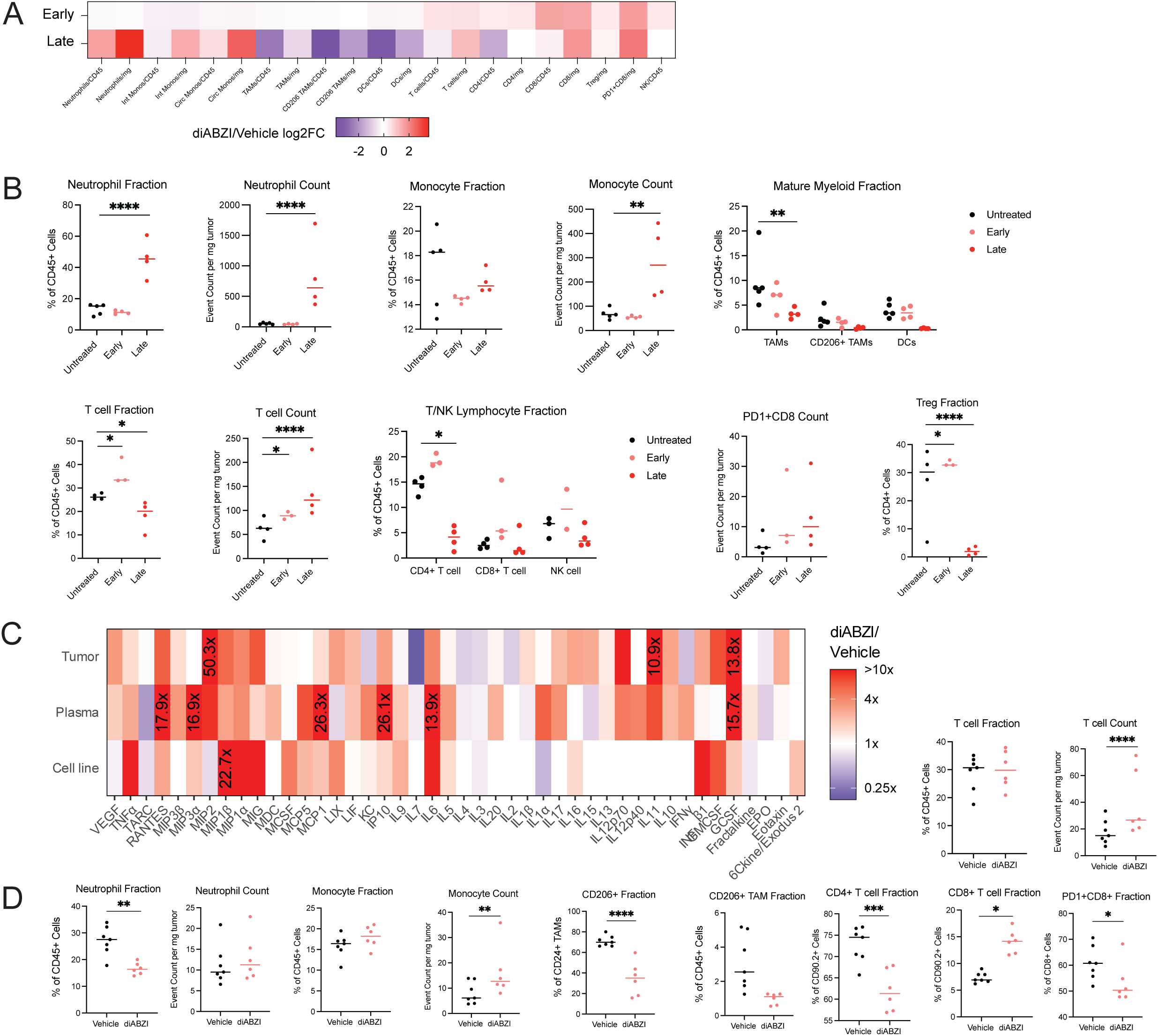
Systemic STING agonism remodels the osteosarcoma tumor microenvironment in vivo. **6A-B.** Analysis of K7M2.1 (BALB/c) OT tumors treated with diABZI (1mg/kg IV) at early (2h) and late (48h) timepoints, comparing vehicle-treated mice to diABZI-treated with gating strategy in Fig. S5, including CD49b NK cell marker. **6A.** Heatmap indicating shifts in immune cell compartments at early and late timepoints after diABZI normalized to untreated tumors, data are shown as fold change (vehicle, n=3 vs. diABZI 2h, n=4; vehicle, n=1 vs. diABZI 48h n=4). **6B.** Original flow cytometry data. Each dot represents data from a single tumor (vehicle, n=4; diABZI 2h, n=4; diABZI 48h n=4, from two independent experiments). **6C.** Cytokine profiling of K7M2.1 cell line conditioned media (vehicle, n=1; diABZI n=1), tumors (vehicle, n=1; diABZI n=1), and plasma (vehicle, n=2; diABZI n=3) 24h after treatment with diABZI. Pro-inflammatory cytokine and chemokine levels (pg/ml) were profiled via two technical replicates for each biological replicate; data are shown as fold change with maximum value as specified. **6D.** Analysis of K7M2.1 tumors at endpoint (18 days post-implantation) after one dose of diABZI, for late timepoint myeloid and lymphoid immunophenotypes. Each dot represents data from a single tumor. Vehicle, n=7; diABZI, n=6. Flow cytometry data are presented as cell type fraction of larger cell type with indicated markers, as well as absolute count normalized to weight of tumor in mg. Data are shown as individual data points with median (B, D) *, P<0.05; **, P<0.01; ***, P<0.001; ****, P<0.0001; two-way ANOVA followed by two-stage step-up multiple comparisons test performed separately for myeloid and lymphoid panels (B, D), non-significant findings are included without p values listed.

We then evaluated cytokine production in K7M2.1 tumor-bearing mice 24h post diABZI treatment. In both plasma and tumors from STING agonist-treated mice, we found elevated levels of pro-inflammatory cytokines and chemokines associated with STING pathway activation, including both CXCL10 and CCL5, cytokines whose corresponding genes were also identified in the STING activation signature (**Fig. 6C**; **Fig 2**). We also observed upregulation of several other cytokines involved in myeloid and T cell recruitment to sites of inflammation, including GCSF, MIP2, CXCL10, CCL2-5, CXCL9, and RANTES. We then profiled K7M2.1 cell line conditioned media to capture the capacity of tumor cells alone to produce cytokines in response to diABZI after 24h. The observed increases in cytokine levels *in vivo* were recapitulated well by conditioned media from STING agonist treated K7M2.1 cell line, suggesting some tumor-intrinsic capacity for cytokine production in response to diABZI (**Fig. 6C**).

To identify remodeling of the tumor microenvironment at later timepoints, K7M2.1 animals were treated on day 8 after implantation and tumors were harvested on day 18 when vehicle tumors reached endpoint (tumor growth curve in **Fig. 5C**). A single dose of diABZI was given in this experiment to enable tumor outgrowth to a degree that allowed immunophenotyping of tumors. Given our finding of infiltration of T cells, neutrophils, and monocytes during the immediate response to diABZI, we sought to test for evolution of these immune cell shifts as tumors continued to grow in the absence of a second, and often tumor-eradicating, dose of diABZI. We observed an absolute increase in monocytes but not neutrophils in these diABZI treated tumors, as well as a decrease in CD206+ expression among TAMs (**Fig. 6D**). In the lymphoid compartment, diABZI-treated tumors evidenced an absolute increase in T cells (when normalized to tumor weight) with a shift from CD4+ to CD8+, as well as a relative decrease in PD1 expression among CD8+ T cells (**Fig. 6D**).

### T cells responses are responsible for anti-tumor efficacy and lasting immunologic memory resulting from STING agonism in vivo

To define whether the diABZI response required cytotoxic CD8+ T cells, diABZI treatment was combined with CD8 T cell depletion using an anti-CD8b antibody (**Fig. 7A-B**). Anti-CD8b treatment did not influence tumor engraftment but did decrease the efficacy of diABZI (**Fig. 7A-B**). After two doses of diABZI, animals with intact CD8 T cells eradicated tumors at a frequency of approximately 50%, whereas only 1 of 8 mice was eradicated a tumor in the CD8b depletion group (**Fig. 7C**).

**Figure 7.**
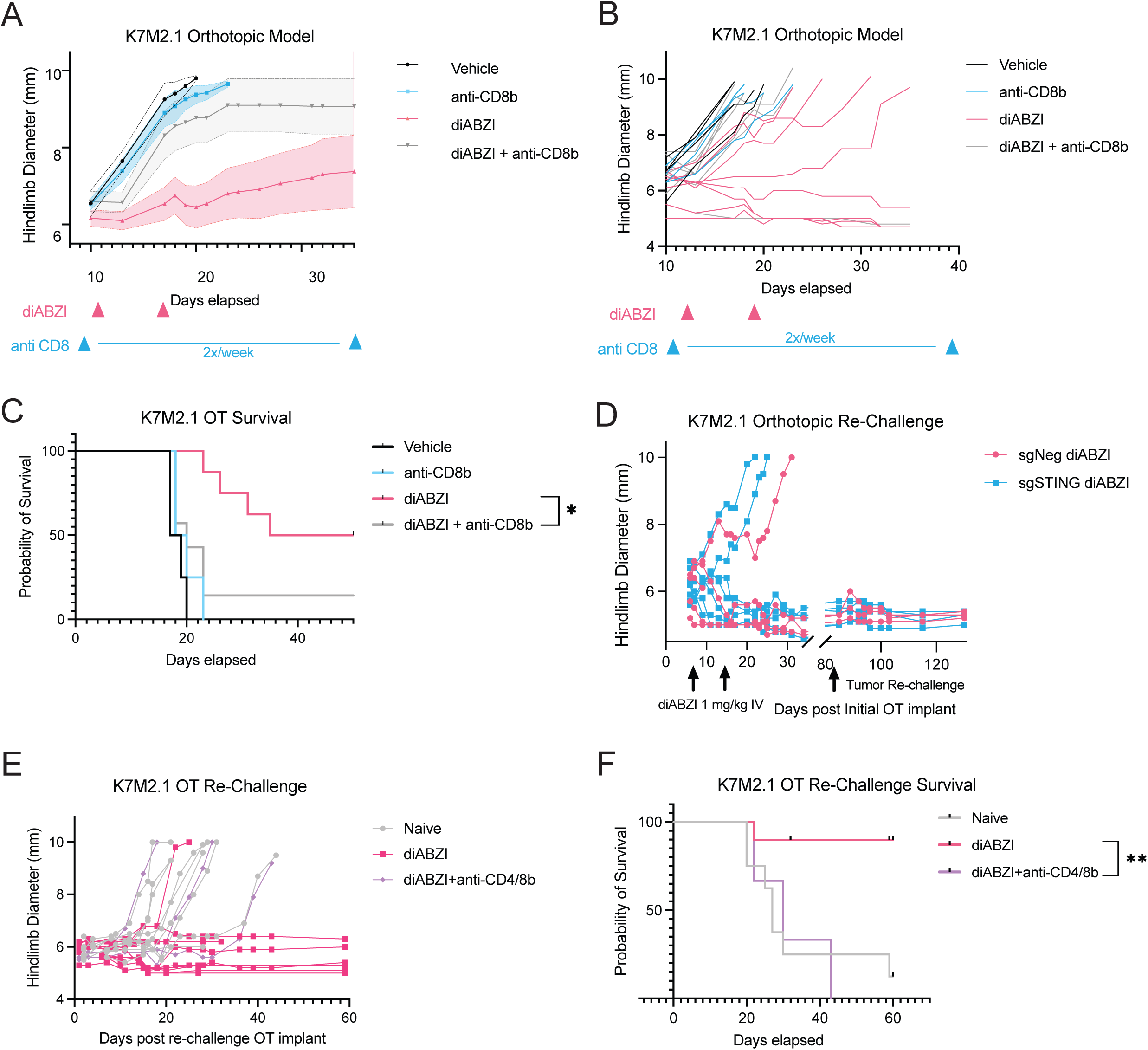
Anti-tumor efficacy and immunologic memory resulting from systemic STING agonism are T cell dependent. **7A-C.** K7M2.1 OT implants in BALB/c mice treated with diABZI (1 mg/kg IV, weekly x 2) and/or CD8b T cell depletion (8mg/kg ip, starting 24h before diABZI, then twice weekly until all tumor-bearing mice reached endpoint), vehicle, n=4; anti-CD8, n=4; diABZI, n=8; anti-CD8b + diABZI, n=7. **7D.** Individual tumor growth curves of diABZI treated mice from Fig. 5D including tumor re-challenge. Mice remaining in study after 80 days were re-challenged with K7M2.1 three months after initial implantation. sgNeg diABZI, n=3; sgSTING diABZI, n=5. **7E-F.** Mice remaining in study after first re-challenge (Fig. 7D) as well as mice from a separate cohort of diABZI-treated animals were subjected to K7M2.1 re-challenge together with tumor naïve control animals, with/without CD4/8 depletion prior to tumor implantation (two dose of twice weekly dosing were given before implantation and continued subsequently). Naïve/no depletion, n=9; Re-challenge/no depletion, n=10;

Since many diABZI-treated mice were tumor-free after two doses of diABZI, we evaluated if this treatment induces lasting immunologic memory by performing tumor re-challenge experiments. When K7M2.1 cells were re-implanted into surviving mice from the K7M2.1 sgNeg/sgSTING OT study (**Fig. 5C**), no tumors were seen two months after implantation, indicating a lasting effect of diABZI treatment in shaping immunological memory (**Fig. 7D**). An additional re-challenge of another cohort of diABZI-treated animals, with and without CD4/8 T cell depletion prior to tumor implantation, demonstrated that the rejection phenotype was abrogated in the absence of T cell-mediated memory (**Fig. 7E-F**). Thus, as expected, long-term memory induced by diABZI is CD4/8 T cell dependent.

## Discussion

There has been widespread interest in studying the cell-intrinsic role of cGAS-STING in cancer as well as the effect of this pathway on shaping the tumor microenvironment ^31,32^. Recent studies suggest that the role of STING activation and IFN signaling can be anti-tumorigenic or pro-tumorigenic in different contexts. This relative balance may depend on cancer-cell intrinsic properties or tumor microenvironment features, calling for close evaluation of the role of STING, including STING targeting therapies, under specific translationally-relevant circumstances ^33^. Here, we report a comprehensive characterization of cGAS-STING in osteosarcoma using human and mouse models (**Graphical Abstract**). We identified STING repression in a subset of patient-derived cell line models and in most osteosarcoma patient samples, in line with prior literature in other genomically unstable cancers ^14,15^. We observed an inverse relationship between chronic cGAS activity and STING repression, which occurs via a post-transcriptional mechanism. Using STING-high cell lines as a resource to evaluate the consequences of STING-dependent inflammation in osteosarcoma patient samples, we defined a “STING activation signature” of the tumor-intrinsic transcriptomic changes in response to STING agonist treatment in osteosarcoma. Enriched pathways involved interferon and inflammatory responses, supporting a tumor-intrinsic interferon signaling capacity in osteosarcoma. Enrichment analysis for this STING signature in treatment-naïve diagnostic biopsies from a bank of sequenced osteosarcoma tumors yielded a range of enrichment across tested samples, supporting a model where the STING signature itself may be localized to stromal and/or immune compartments. While the STING activation signature was experimentally derived by treating tumor cell lines with a STING agonist, its enrichment in patient samples may reflect transcriptional changes in the tumor microenvironment of patient samples due to other upstream stimulators of type 1 IFN production from distinct DNA or RNA sensing pathways. High enrichment with the STING activation signature was protective on overall and disease-free survival independent of metastatic status at presentation. Considering other literature that nominates increased immune infiltration as a favorable prognostic feature in osteosarcoma, our work directly implicates STING activation as a specific inflammatory pathway that may be involved in establishing or maintaining this tumor microenvironment feature ^4^.

Single cell analysis revealed that the “STING activation” signature was only found in a small subset of tumor cells, which may be due to repression of either cGAS or STING that we have observed (**Fig. 1; Fig. S1**) and may allow osteosarcoma to evade inflammatory response to cytosolic DNA that results from ongoing CIN and/or chemotherapy. Multiple additional cell types in the osteosarcoma tumor microenvironment evidence inflammatory signaling as captured by this STING activation signature, including non-tumor stromal cells, immune cells (both myeloid and lymphoid), endothelial cells, and osteoblasts. Endothelial cell STING has been described to crosstalk with tumor cGAMP and enhance the efficacy of combinatorial immunotherapy for liver cancer ^34^. While tumor endothelial cells facilitate osteosarcoma pathogenesis, and targeting of angiogenesis is being studied in the current osteosarcoma clinical trial AOST2032, the potential role of endothelial cells in mediating immunotherapy response in osteosarcoma has not been adequately considered. The STING inferred signature is also enriched in osteoclasts in the treatment-naïve samples, which is intriguing given prior work indicating that STING-mediated tonic expression of type 1 interferons maintains osteoclasts in a precursor, de-differentiated state ^35^. Thus, inflammatory signaling in osteoclasts may affect osteoclast cell state, which in turn could affect osteoclast contributions to osteosarcoma pathogenesis.

Spatial transcriptomics performed on metastatic osteosarcoma specimens demonstrated that the tumor core was largely devoid of inflammatory signal across all specimens analyzed, indicating suppression of inflammatory signaling in advanced metastatic disease. These metastatic osteosarcomas are indeed “immune excluded,” characterized by inflammatory signaling and immune cell infiltration only in the peritumoral area, akin to a foreign body response ^36^. The absence of T cell infiltration in these tumors supports a rationale for the utilization of a STING agonist, which has been shown here to initiate a cytokine program that increases T cell infiltration into osteosarcoma lesions.

To corroborate the findings of tumor STING repression in patient-derived models, immunocompetent osteosarcoma murine models were employed to directly probe the effects of tumor STING on tumor behavior *in vivo*. In both syngeneic cell line models tested, tumor STING loss did not alter tumor growth, metastatic propensity, or tumor microenvironment constitution, consistent with prior literature indicating that perturbation of STING in cell lines in a transplantation setting is not sufficient to modulate tumor behavior ^32,37,38^. Other work has supported a pro-metastatic role for STING in animal models ^17^. Considering the STING repression observed in both patient-derived models and osteosarcoma tumors, it remains plausible that STING repression impacts tumor behavior in the context of early tumor evolution, which may not be recapitulated in these models. Future studies of the role of tumor and host STING KO in a genetically engineered mouse model of osteosarcoma may elucidate the impact of STING repression in the critical context of tumor evolution. Given the protective effect of STING activation in patients, we sought to evaluate the translational potential of STING agonism with diABZI, a unique agent in contrast to virtually all other STING agonists which require intratumoral administration ^19^. In syngeneic OT models of osteosarcoma, a single dose of diABZI restrained tumor growth and altered late immune infiltrate in the K7M2.1/BALB/c model and two doses of diABZI improved survival in the highly aggressive F420/B6 model, highlighting the relevance of this therapy across murine models reflecting different immune skews. Importantly, in the K7M2.1/BALB/c model, two weekly doses of diABZI were sufficient to allow for long term survival of mice regardless of the presence or absence of tumor STING. This result supports a primary role for host STING in the context of therapy-induced STING activation ^32,37,39^. Consistent with these *in vivo* findings, a tumor-macrophage co-culture assay revealed the relative greater contribution of myeloid STING, compared to tumor STING, in the production of IFN-β in response to STING agonism.

Evidence of IFN-β induction in STING agonist-treated tumors suggests that anti-tumor and curative responses occur through type 1 interferon signaling, corroborated by a robust induction of pro-inflammatory cytokines observed within 24h after treatment. GCSF and MIP2 likely contribute to the recruitment of neutrophils to the tumor microenvironment that we observed by flow cytometry. In STING-agonist treated samples compared to vehicle-treated samples, we found elevated levels of many cytokines and chemokines that recruit monocytes to sites of inflammation, including CXCL10 and CCL2-4, likely responsible for observed monocyte recruitment to tumors at a 48h timepoint. In addition, CXCL9 and CCL5/RANTES were induced in diABZI-treated samples and have been described to mediate recruitment of Th1 CD4+ cells and CD8+ cytotoxic T cells to areas of inflammation.

Flow cytometric profiling of early and late shifts in tumor microenvironment constitution in the K7M2.1/BALBc model further clarified the time course of immune cell recruitment to the orthotopic tumor microenvironment. We concluded from these findings that STING activation in the tumor microenvironment results in T cell recruitment to the tumor followed by acute inflammation associated with monocyte and neutrophil infiltration. This rapid infiltration of CD8 T cells by 2 hours suggests that these CD8 T cells have previously been activated by tumor antigen likely following orthotopic implantation/during early tumorigenesis in this model, and that diABZI increases cytokine production and CD8 T cell homing to the orthotopic tumor microenvironment. We also saw a shift in TIL proportions away from mature myeloid cells, such as TAMs and DCs, likely due to the increase in peripherally recruited monocytes and neutrophils, in addition to T cells. Direct cytotoxic effects of STING agonism on the DC population have been described previously ^40^, and previous literature has also shown transient accumulation of a CD11b^mid^Ly6c+F480+ monocyte population in multiple syngeneic tumor models after cGAMP administration ^39^.

We identified that the long-term efficacy of systemic diABZI is CD8+ T cell dependent, consistent with prior reports studying intratumoral cGAMP administration, though the magnitude of effect appears to be greater with systemic administration in this disease context ^39,41^. A tumor re-challenge experiment of long-term surviving mice previously treated with two doses of diABZI yielded rejection in 100% of mice, supporting the presence of long-term immunologic memory resulting from diABZI monotherapy. We also demonstrated that this rejection requires CD4/8 T cells, supporting a central role for the T cell compartment in both immediate and long-term immune remodeling that is initiated by STING agonism.

This work defines the spectrum of STING function and repression across a diverse array of osteosarcoma models representing this highly aggressive pediatric cancer and investigates the therapeutic potential of STING agonism while critically delineating that tumor STING is not required to drive anti-tumor responses. Currently, a non-cyclic dinucleotide STING agonist is being studied alone or in combination with immune checkpoint blockade in adults with relapsed/refractory solid tumors (NCT03843359). This preclinical work nominates STING agonism as a novel modality to achieve beneficial innate immune activation in patients despite repression of the cGAS-STING pathway in osteosarcoma tumors and calls for further discovery related to immunotherapy approaches for this highly aggressive pediatric cancer.

## Methods

### Sex as a biological variable

Our study examined male and female animals, and similar findings are reported for both sexes. Due to observed variability in some cell line growth rates in males vs. females, the drug study shown in **Fig. 5B-C** examined female B6 mice to facilitate treatment initiation at a similar tumor size across the cohort; findings are expected to be relevant for males.

### Cell lines and mouse strains

Human osteosarcoma patient-derived tumor xenograft-derived cell lines OS052, OS152, OS186, OS384, OS457, OS525, OS526, OS742, OS766, OS774, and OS833 were generated as described previously ^42^. Human OS cell lines 143B, SJSA-1, MG-63, Saos-2, U-2OS, murine OS cell line K7M2, and human fibroblast cell line IMR-90 were obtained from the American Type Culture Collection (ATCC). Murine OS cell line F420 was gifted by Dr. Jason Yustein. K7M2.1 was derived from an orthotopic K7M2 tumor (in BALB/c), chopped into small fragments and digested per standard protocol (see below). All cell lines were maintained in DMEM medium (Gibco, 11-965-092) supplemented with 10% bovine growth serum (Hyclone, SH30541.03) and 1% penicillin-streptomycin (Corning, MT30002CI), unless otherwise stated. Bone marrow-derived macrophages were maintained in αMEM medium (Gibco, 12561056) supplemented with 10% fetal bovine serum (Corning, #35-011-CV) and 1% penicillin-streptomycin. All cells were maintained at 37°C under 5% CO2 and were regularly tested for STR analysis and mycoplasma contamination. BALB/cJ (strain #:000651), C57BL/6J (strain #:000664), and B6(Cg)-*Sting1^tm^*^1^*^.2Camb^*/J (strain #:025805) were obtained from The Jackson Laboratory.

Murine osteosarcoma cell lines (K7M2.1, F420) with *Sting1* KO were generated by Cas9 ribonucleoprotein nucleofection using a Lonza 4D-Nucleofector and SE Pulse Code DS-150 Cell Line Kit. Three guides were used simultaneously (GCGAGGCUAGGUGAAGUGCU; GAUGAUCCUUUGGGUGGCAA; ACCUGCAUCCAGCCAUCCCA). Based on KO efficiency by immunoblotting, a ratio of sgRNA to Cas9 of 9:1 was used for K7M2.1 and a ratio of 4.5:1 was for F420.

### Patient sample collection

Patients aged 0-39 with a diagnosis of osteosarcoma were included as candidates for this study. Samples were collected at Lucile Packard Children’s Hospital (n=9), UCSF Benioff Children’s Hospital, San Francisco (n=13) and Oakland (n=1). Informed consent was obtained for all patients and studies were approved by local IRBs. Germline DNA was obtained from peripheral blood. Tumor samples were either obtained from FFPE blocks or collected fresh, snap frozen in LN2, and embedded in OCT. 5 µm cryosections were stained with H&E and evaluated by a pediatric pathologist. Regions estimated to have ≥70% tumor were macrodissected, disrupted using a mortar and pestle under LN2 conditions, and homogenized with a QIAshredder (QIAgen, 80204).

### IHC, immunofluorescence, and Visium spatial profiling

FFPE sections were deparaffinized in xylene and rehydrated using an ethanol series and water. Antigen retrieval was achieved by incubating slides in citrate buffer pH 6.0 for 10min at 100°C in a pressurized chamber. Slides were blocked in 2.5% normal horse serum (Vector Laboratories, S-2012-50), followed by overnight incubation with primary antibody at 4°C and secondary antibody conjugated to a horseradish peroxidase polymer (Vector Laboratories, MP-7401). Images were captured on an AxioScan 7 at 20X magnification (Zeiss). For quantitative analysis of IHC data, the Cytonuclear Module v2.0.9 of HALO v3.6.4134.362 (Indica Labs) was utilized to determine labeling intensity in patient tissues. Results for each individual patient was visualized on a heatmap using Morpheus (https://software.broadinstitute.org/morpheus).

For immunofluorescence, cells were fixed with formalin for 10min, permeabilized using 0.1% Triton-X in PBS for 10min on ice, and blocked in 5% species-specific serum in PBS, followed by overnight incubation with primary antibody at 4°C. Secondary antibody incubation occurred for 1h. Cells were mounted with mounting media containing DAPI. Slides were imaged on a Leica microscope and the percentage of micronuclei (MN) per nuclei, in addition to co-positivity for cGAS, were manually quantified for at least 5 fields of view (FOV) per cell line using a 20X objective.

Visium spatial profiling was performed as described previously^36^. The CytoSig (10.1038/s41592-021-01274-5) package was used to infer type I interferon signaling activity ^26^. The “STING activation signature” score was quantified using the AddModuleScore_UCell function in the UCell package (<u>10.1016/j.csbj.2021.06.043</u>). Spatial maps of the CytoSig type I interferon activity score and the STING activation signature score were generated using the SpatialFeaturePlot function in Seurat and plotted on different scales.

### In vitro drug treatments and cGAMP quantification

Cells were treated with diABZI (5 µM), G3-YSD (1 µg/mL), bafilomycin (200 nM), or G150 (1 µg/mL) prepared according to manufacturer’s instructions and collected at 6h and 24h for immunoblotting and RT-qPCR/RNAseq respectively, unless otherwise stated. cGAMP was quantified in cell lysates as described previously ^17^, with the following modification: whole-cell lysates were generated by lysing the cell pellet in RIPA (Sigma-Aldrich, #R0278) with 5 mM EDTA. cGAMP ELISA was performed according to the manufacturer’s instructions (DetectX Direct 2’-3’-Cyclic GAMP Enzyme Immunoassay Kit, Arbor Assays, #K067-H1).

### Immunoblotting

Whole-cell lysates were prepared in RIPA (Sigma-Aldrich, #R0278) containing protease and phosphatase inhibitors (Pierce, #A32953, #A32957). Protein concentrations were determined by Pierce BCA Protein Assay Kits (Thermo Fisher, #23227). Samples were mixed with Laemmli SDS sample buffer (Alfa Aesar, #J60015), denatured at 95°C x 5min, electrophoresed via SDS-PAGE in 4-15% Mini-PROTEAN TGX Precast Protein gels (Bio Rad, #4561086) and transferred onto PVDF membranes using Trans-Blot Turbo (Bio Rad). Membranes were blocked with 5% milk in TBST (Bioland Scientific, #TBST01-03) at room temperature (RT), then incubated with primary antibodies in 5% BSA (Sigma Aldrich, #A4503) overnight at 4°C, followed by secondary antibody incubation at RT. See Supplemental Table 4 for primary antibody information. Immunoblots were incubated with ECL (Bio Rad, #170-5061) and visualized using an Odyssey fluorescence scanner (Li-Cor). Quantification was performed using Li-Cor Image Quantification Software.

### Nucleic acid extraction, library preparation, and RT-qPCR

Tumor DNA and RNA were co-extracted from patient samples using the AllPrep kit (QIAgen, 80204) or the AllPrep FFPE kit (QIAgen, 80234). RNA was extracted from osteosarcoma PDX cell lines using the RNeasy Kit (Qiagen) according to the manufacturer’s instructions with a DNAse I digestion. Germline DNA was extracted from peripheral blood with the DNeasy Blood and Tissue kit (QIAgen, 69504). DNA was quantified on the Nanodrop 2000 (Thermo Fisher) and QuBit HS DNA assay (Thermo Fisher, Q32851). DNA integrity was quantified with Genomic ScreenTape (Agilent, 5067-5365) on the TapeStation 4200 (Agilent) or the HS-gDNA kit (Agilent, DNF-468-0500) on the Fragment Analyzer (Agilent). RNA was quantified on the Nanodrop 2000. RNA integrity number and DV200 were evaluated using the HS RNA ScreenTape (Agilent, 5067-5579) and the HS-RNA kit (Agilent, DNF-472-0500) Fragment Analyzer (Agilent).

WGS libraries were made using the TruSeq Nano kit (Illumina, FC-121-4001) with a 350bp insert as per manufacturer’s instructions using 100ng of germline DNA and sequenced to a depth of 30x, or an input of 200ng of tumor DNA and sequenced to a depth of 60x. Libraries were assessed with the D1000 ScreenTape (Agilent, 5067-5062) to determine library size and distribution, and absence of primer dimers. Libraries were sequenced on the HiSeq X Ten or NovaSeq 6000 at 2×150bp. RNAseq libraries were made using the TruSeq Stranded mRNA kit (Illumina, RS-122-2101) or the RNA Exome kit (Illumina, 20020189) per the manufacturer’s instructions. All manufacturer’s controls were used in preparation of the libraries. Libraries were assessed with the HS NGS (Agilent, DNF-474) or the HS D1000 (Agilent, 5067-5585) assays to determine library size and distribution, and the absence of primer dimers. Sequencing was performed on the following instruments: HiSeq 4000 2×75, 2×100; CHiSeq 4000 2×100, 2×150, NovaSeq 2×150; HiSeq Xten 2×150. Bulk RNAseq was performed on OS PDX derived cell lines as described previously ^18^.

cDNA synthesis was performed with 1μg RNA using Maxima First Strand cDNA Synthesis Kit for RT-qPCR (Thermo Fisher, #K1641). RT-qPCR was performed using 25ng cDNA with SYBR Green PCR Master Mix using a thermocycler (Biorad). Data were normalized to Actin or GAPDH for human or mouse respectively. Fold induction was calculated by ΔΔ*C*_t._ Gene specific primers include: ISG15 (GAACTCATCTTTGCCAGTA, ATCTTCTGGGTGATCTGC), CXCL10 (TACCTGCATCAGCATTAGTA, TGTAGCAATGATCTCAACAC), hSTING (CACTTGGATGCTTGCCCTC, GCCACGTTGAAATTCCCTTTTT; AGCATTACAACAACCTGCTACG, GTTGGGGTCAGCCATACTCAG), GAPDH (AGGTCGGTGTGAACGGATTTG, TGTAGACCATGTAGTTGAGGTCA), Actin (CATTCCAAATATGAGATGCGTTGT, GCTATCACCTCCCCTGTGTG).

### Quantification of cytokines

Cell culture supernatant was collected at 24h post-stimulation, filtered through 0.45μm, and centrifuged at 4°C, 3000 x g, x 10min. For plasma and tumor analysis, at 24h post-STING agonist treatment, whole blood and tumors were collected from K7M2.1 tumor-bearing mice. Blood was collected in EDTA tubes and centrifuged at 1000 x g for 10 min at 4°C. Plasma samples were diluted 3-fold in PBS. Tumors were homogenized in cold RIPA buffer containing Sigma protease inhibitors. Tissue homogenate was centrifuged at 2000 x g for 15min at 4°C. Protein quantification and normalization were performed by Eve Technologies (Calgary, Alberta). Samples were subjected to the Human Cytokine/Chemokine Panel A 48-Plex Discovery Assay® Array (HD48A) or the Mouse Cytokine/Chemokine 44-Plex Discovery Assay® Array (MD44) using the Luminex^TM^ 200 system (Luminex, Austin, TX, USA) by Eve Technologies (Calgary, Alberta). Results were reported as observed concentration estimated by a standard curve as per the manufacturer’s instructions.

### BMDM isolation, differentiation, and profiling

Bone marrow-derived macrophages (BMDMs) were obtained from C57BL/6J or B6(Cg)-*Sting1^tm^*^1^*^.2Camb^*/J mice and isolated and differentiated as described previously ^43^, with the exception of using 10% CMG14-12 conditioned DMEM as a source of M-CSF ^44^. To assess the production of IFN-β, cells were plated in isolation and in co-culture (at a ratio of 1:1) and treated with 5uM diABZI. Conditioned media was collected at 24h post-stimulation, filtered through 0.45μm, and centrifuged at 4°C, 3000 x g, x 10min. IFN-β production was detected with Verikine Mouse IFN-β ELISA (PBL, #42400-1) according to the manufacturer’s instructions.

### In vivo studies

For OT studies, paratibial tumor cell line implantation was employed as described previously ^18^. Tumor growth was monitored with twice-weekly measurements of the lower hindlimb until the tibial diameter reached 1 cm (primary tumor endpoint). Metastasis study endpoint following hindlimb amputation occurred when mice were evaluated to have significant respiratory distress requiring euthanasia or when lung burden had increased >20% by microCT. For IV metastasis studies, 1×10^6^ osteosarcoma cells in PBS were injected via the lateral tail vein per standard procedure and mice were euthanized at a standard timepoint. For metastatic studies, lungs were harvested for histological processing ^18^. H&E slide scans were manually analyzed for metastatic burden using Fiji. Tumor burden (% Area) was calculated as percentage of tumor area per lung area.

For drug studies, cells were implanted OT as described above. Mice were randomized to treatment with vehicle control or diABZI IV via retro-orbital injection. For drug preparation, diABZI stock (Selleck Chem, #S8796) was prepared according to the manufacturer’s instructions (5% DMSO + 40% PEG300 + 5% Tween 80 + 50% ddH2O). Weight was monitored daily for three days following each diABZI dose and otherwise weekly. Supportive care with diet gel and forage mix were given to all study animals, and any animals with >5% weight loss were given carprofen at 0.5 mg/kg/dose. For T cell depletion, anti-CD4 mAb (GK1.5; Bio X Cell, #BE0003) and/or anti-CD8b mAb (53-5.8; Bio X Cell, #BE0223) was prepared at 8mg/kg/dose according to manufacturer’s instructions and given IP twice weekly.

### Tumor Digestion

Freshly isolated primary tumors were processed same day or minced and viably frozen in 90%FBS/10%DMSO. Tumors were digested in 5mL HBSS + 5% FBS + 0.1% HEPES supplemented with 7.5mg Collagenase and 50µL TURBO DNAse (Invitrogen) and incubated at 37°C on a shaker x 1.5h. Tubes were vortexed on low speed every 15-20min during incubation. Once the tumor tissue was thoroughly digested, 20 mL buffer (cold PBS + 2% FBS) was added to each sample. Samples were vortexed for 30sec on high speed and contents were passed through a 70µm filter. Single-cell suspensions were spun at 4°C, 500 x g, x 5min and supernatant was decanted. Cells were resuspended in 2mL ACK Lysing Buffer (Gibco, #A10492-01) for 3min and quenched with buffer. Spin was repeated and cells were resuspended in cold bbuffer on ice.

### Immunophenotyping

Tumors were immunophenotyped via multiparametric flow cytometry using myeloid panel and/or T-lymphoid panels. Spins were performed at 4°C, 500 x g, x 5min. 0.5–1 x 10^6^ live cells/sample were stained for each panel in a V-bottom 96-well plate. Cells were stained for viability using the LIVE/DEAD Fixable Aqua Dead Cell Stain Kit (Life Technologies, #L34966) with a 20min incubation on ice in the dark. After a wash, cells were resuspended in 50µL antibody stain master mix/1 x 10^6^ live cells and stained for 30 min on ice in the dark. Master mix included antibodies, mouse Fc block 2.4G2 (Anti-CD16 + CD32 Rat Monoclonal Antibody, Tonbo Biosciences, #10051), Brilliant Stain Buffer (BD Biosciences, #563794), and PBS + 2% FBS. See Supplemental Table 5 for antibodies used in myeloid and lymphoid panels. Single color compensation controls were prepared using UltraComp beads (Invitrogen, #01-2222-41). Samples were analyzed using the BD FACSAria Fusion Flow Cytometer (PFCC, UCSF) and data was analyzed using FlowJo software.

### Genomic analysis

Whole genome analysis, including purity estimation, was performed as described previously ^42^. For gene expression analysis, raw FASTQ files were quality checked (QC) with FastQC and adapter trimmed with Trimmomatic_0.36 using paired end mode ^45^. Reads were mapped to the hg38 human genome (GENCODE p5) with STAR_2.5.3 aligner ^46^. Gene level counts were calculated with STAR–*quantMode* option. QC alignments were further assessed with ngsutilsj_0.4.16. Counts per million (CPM) were calculated after normalization using the trimmed mean of M-values from raw counts, followed by log2 transformation with prior.count set to 2 using the EdgeR_3.28.1 package ^47^. Differential expression analysis was performed using the VOOM method followed by linear modeling and empirical Bayes moderation with the limma (v3.42.2) package to obtain p-values, adjusted p-values and log-fold changes (logFC) ^48^. Gene Set Enrichment Analysis was performed with ranked logFC using the R package fgsea_1.15.1. Hallmark gene sets and KEGG pathways were downloaded from Molecular Signatures Database (MSigDB).

Estimation of immune cell components in RNAseq data was calculated using CIBERSORTX ^49^. Sample purity was determined based upon manual curation of WGS using allele-specific copy-number and variant allele frequencies (VAF). Clonal fraction was estimated as percentage of single nucleotide variants that were present at an VAF > 0.8. TMB was calculated as the variant burden across the whole genome (in variants/Mb). The percentage of microsatellites with repeat instability was calculated using msisensor, where values greater than 2-3% is considered microsatellite instable ^50^. Total HLA I, B2M, PD-L1, B7, and CD47 expression was calculated from the RNAseq data. The immune score and stromal score were calculated using the ESTIMATE package for R ^51^. Samples were k-means clustered into 4 groups based upon the CIBERSORTX scores for the 22 immune cell types included in the LM22 scoring matrix ^52^.

### Single cell RNAseq analysis

Raw FASTQ files were obtained from two publicly available datasets, GSE152048 and GSE162454 and independently processed using the same analytical pipeline. Alignment was performed using Cell Ranger 10x Genomics, v7 (aligned to the hg38 reference genome), with downstream analysis in R. Filtered data matrices were imported into R and processed using Seurat (v5.0). For each sample, cells were filtered based on number of genes (>200 and <4000), percentage of mitochondrial reads (<15%), and total RNA count (nCount_RNA>1000). Normalization was completed using SCTransform regressing for percent mitochondria. Samples were merged and dimensionality reduction was performed using Principal Component Analysis (PCA), retaining the top 50 principal components. Uniform Manifold Approximation and Projection (UMAP) was applied to the first 30 principal components for visualization. Cell clustering was performed using Seurat FindNeighbors and FindClusters functions (resolution parameter of 1.2) utilizing the same 30 principal components. Resulting clusters were visualized on the UMAP plot to delineate distinct cell populations. Harmony (v1.1) was employed to mitigate batch effects across samples. Post-integration, PCA (30 components), UMAP embeddings, and cell clusters were recomputed using the Harmony-corrected embeddings, and clustering was performed with a resolution of 1.0. Cell type annotations were determined via a manual curation process involving analysis of positive and negative marker gene expression and considering both expression levels and percentage of cells expressing each marker (Supplemental Table 3). Overall structure, clustering patterns, and proximity of clusters in the UMAP visualization were considered as supporting evidence for cell type assignments. ScType was employed to independently validate the manually curated cell groups, and this tool was leveraged to assess the prevalence of specific gene signatures in each dataset. Single Cell Variational Aneuploidy Analysis (SCEVAN) was applied to distinguish between malignant and non-malignant cells.

### Survival analysis

Survival analysis was conducted on a cohort of treatment-naïve patients with primary osteosarcoma (n=23). External validation was performed with an independent osteosarcoma dataset (n=72) obtained from the UCSC Toil RNA-seq pipeline (https://github.com/BD2KGenomics/toil-rnaseq), with corresponding clinical metadata retrieved from UCSC Xena. Gene signatures for STING activation and STING inferred were used to stratify sample employing a median split approach, with samples above the 50th percentile classified as ‘high’ and those below as ‘low’ for each signature. Survival analyses were performed using the ‘survival’ package (version 3.1.11) in R (see below). Cox proportional hazards models were fitted to estimate hazard ratios and their corresponding 95% confidence intervals. To ensure harmonization between cohorts, the UCSC Toil RNA-seq pipeline was used to produce Transcripts Per Million (TPM) values, which served as the input for these analyses ^53^.

### Statistics

Data were analyzed using GraphPad Prism 10 (GraphPad Software). Comparisons between 2 groups were analyzed using a 2-tailed unpaired Student’s t test. Comparison between multiple groups were analyzed using 1-way ANOVA with repeated measures or 2-way ANOVA with two-stage step up multiple comparisons test. For survival analyses, survival curves were estimated for each group using the Kaplan-Meiser method and compared statistically using the log-rank test. Statistical significance was defined as a P value of less than 0.05.

### Study approval

For patient sample collection, informed consent was obtained for all patients and studies were approved by local IRBs. All animal studies were approved by IACUC at University of California San Francisco.

### Data availability

The Supporting Data Values file includes underlying graphed data presented in both the main text and supplemental material. The cell line data shown in Fig. 2 and Fig. S2 are available in GEO under accession GSE275335. Patient data shown in Fig. 2 are available in dbGaP under dbGaP accession phs002430. The data shown in Fig. 3 and Fig. S3 are available in GEO under accession GSE152048 and GSE162454. The Visium spatial transcriptomics data shown in Fig. 4 is available as described in the published manuscript ^36^ and raw data is available on request.

## Supporting information

All Supplemental Information

## Acknowledgments

EASC was funded by the National Cancer Institute (R01CA243555) and by a grant from the Osteosarcoma Institute (OSI). EPY was funded by grants from St. Baldrick’s Foundation, Battle Osteosarcoma, Alex’s Lemonade Stand Foundation, MIB Agents, Hyundai Hope on Wheels, and CURE Childhood Cancer. AGL was funded by the National Cancer Institute (R50CA274213). CRS was funded by a grant from the Rally Foundation for Childhood Cancer Research. This research was supported in part by the Intramural Research Program of the National Institutes of Health (NIH). The contributions of the NIH author(s) were made as part of their official duties as NIH federal employees, are in compliance with agency policy requirements, and are considered Works of the United States Government. However, the findings and conclusions presented in this paper are those of the author(s) and do not necessarily reflect the views of the NIH or the U.S. Department of Health and Human Services. This work was also supported by the University of California, San Francisco Preclinical Therapeutics Core (PTC) under NIH/NCI awards P30CA082103 and S10OD025022. We acknowledge the PFCC (RRID:*SCR_018206*) supported in part by Grant NIH P30 DK063720 and by the NIH S10 Instrumentation Grant S10 1S10OD021822-01. Sequencing was performed in part at the UCSF CAT, supported by UCSF PBBR, RRP IMIA, and NIH 1S10OD028511-01 grants. We thank Dr. Jason Yustein (Emory University) for the kind gift of the F420 cell line. We thank Sarah Pyle for graphic design and Andrew Clugston for help with data visualization. We thank all members of the Sweet-Cordero laboratory for helpful suggestions and critiques. We are grateful to all the patients and their families who generously allowed us to utilize tumor and blood samples for research by signing informed consent.

## References

1 Meltzer, P. S. & Helman, L. J. New Horizons in the Treatment of Osteosarcoma. N Engl J Med 385, 2066–2076 (2021).

2 Chen, X. et al. Recurrent somatic structural variations contribute to tumorigenesis in pediatric osteosarcoma. Cell reports 7, 104–112 (2014).

3 Behjati, S. et al. Recurrent mutation of IGF signalling genes and distinct patterns of genomic rearrangement in osteosarcoma. Nat Commun 8, 15936 (2017).

4 Cillo, A. R. et al. Ewing Sarcoma and Osteosarcoma Have Distinct Immune Signatures and Intercellular Communication Networks. Clin Cancer Res 28, 4968–4982 (2022).

5 Wu, C. C. et al. Immuno-genomic landscape of osteosarcoma. Nat Commun 11, 1008 (2020).

6 Tawbi, H. A. et al. Pembrolizumab in advanced soft-tissue sarcoma and bone sarcoma (SARC028): a multicentre, two-cohort, single-arm, open-label, phase 2 trial. Lancet Oncol 18, 1493–1501 (2017).

7 Le Cesne, A. et al. Programmed cell death 1 (PD-1) targeting in patients with advanced osteosarcomas: results from the PEMBROSARC study. Eur J Cancer 119, 151–157 (2019).

8 Beird, H. C. et al. Osteosarcoma. Nat Rev Dis Primers 8, 77 (2022).

9 Mackenzie, K. J. et al. cGAS surveillance of micronuclei links genome instability to innate immunity. Nature 548, 461–465 (2017).

10 Samson, N. & Ablasser, A. The cGAS-STING pathway and cancer. Nat Cancer 3, 1452–1463 (2022).

11 Bakhoum, S. F. & Cantley, L. C. The Multifaceted Role of Chromosomal Instability in Cancer and Its Microenvironment. Cell 174, 1347–1360 (2018).

12 Kwart, D. et al. Cancer cell-derived type I interferons instruct tumor monocyte polarization. Cell reports 41, 111769 (2022).

13 Konno, H. et al. Suppression of STING signaling through epigenetic silencing and missense mutation impedes DNA damage mediated cytokine production. Oncogene 37, 2037–2051 (2018).

14 Falahat, R. et al. Epigenetic reprogramming of tumor cell-intrinsic STING function sculpts antigenicity and T cell recognition of melanoma. Proc Natl Acad Sci U S A 118 (2021).

15 Low, J. T. et al. Epigenetic STING silencing is developmentally conserved in gliomas and can be rescued by methyltransferase inhibition. Cancer Cell 40, 439–440 (2022).

16 Song, S. et al. Decreased expression of STING predicts poor prognosis in patients with gastric cancer. Sci Rep 7, 39858 (2017).

17 Li, J. et al. Non-cell-autonomous cancer progression from chromosomal instability. Nature 620, 1080–1088 (2023).

18 Schott, C. R. et al. Osteosarcoma PDX-Derived Cell Line Models for Preclinical Drug Evaluation Demonstrate Metastasis Inhibition by Dinaciclib through a Genome-Targeted Approach. Clinical Cancer Research 30, 849–864 (2024).

19 Ramanjulu, J. M. et al. Design of amidobenzimidazole STING receptor agonists with systemic activity. Nature 564, 439–443 (2018).

20 Barber, G. N. STING: infection, inflammation and cancer. Nat Rev Immunol 15, 760–770 (2015).

21 Gonugunta, V. K. et al. Trafficking-Mediated STING Degradation Requires Sorting to Acidified Endolysosomes and Can Be Targeted to Enhance Anti-tumor Response. Cell reports 21, 3234–3242 (2017).

22 Jeltema, D. et al. STING trafficking as a new dimension of immune signaling. J Exp Med 220 (2023).

23 Mannheimer, J. D. et al. Transcriptional profiling of canine osteosarcoma identifies prognostic gene expression signatures with translational value for humans. Commun Biol 6, 856 (2023).

24 Zhou, Y. et al. Single-cell RNA landscape of intratumoral heterogeneity and immunosuppressive microenvironment in advanced osteosarcoma. Nat Commun 11, 6322 (2020).

25 Liu, Y. et al. Single-Cell Transcriptomics Reveals the Complexity of the Tumor Microenvironment of Treatment-Naive Osteosarcoma. Front Oncol 11, 709210 (2021).

26 Jiang, P. et al. Systematic investigation of cytokine signaling activity at the tissue and single-cell levels. Nat Methods 18, 1181–1191 (2021).

27 Khanna, C. et al. An orthotopic model of murine osteosarcoma with clonally related variants differing in pulmonary metastatic potential. Clin Exp Metastasis 18, 261–271 (2000).

28 Zhao, S. et al. NKD2, a negative regulator of Wnt signaling, suppresses tumor growth and metastasis in osteosarcoma. Oncogene 34, 5069–5079 (2015).

29 Bakhoum, S. F. et al. Chromosomal instability drives metastasis through a cytosolic DNA response. Nature 553, 467–472 (2018).

30 Beltra, J. C. et al. Developmental Relationships of Four Exhausted CD8(+) T Cell Subsets Reveals Underlying Transcriptional and Epigenetic Landscape Control Mechanisms. Immunity 52, 825–841 e828 (2020).

31 Tian, Z. et al. Cancer immunotherapy strategies that target the cGAS-STING pathway. Frontiers in Immunology 13 (2022).

32 Schadt, L. et al. Cancer-Cell-Intrinsic cGAS Expression Mediates Tumor Immunogenicity. Cell reports 29, 1236–1248 e1237 (2019).

33 Lanng, K. R. B. et al. The balance of STING signaling orchestrates immunity in cancer. Nat Immunol (2024).

34 Lv, H. et al. TET2-mediated tumor cGAS triggers endothelial STING activation to regulate vasculature remodeling and anti-tumor immunity in liver cancer. Nat Commun 15, 6 (2024).

35 MacLauchlan, S. et al. STING-dependent interferon signatures restrict osteoclast differentiation and bone loss in mice. Proc Natl Acad Sci U S A 120, e2210409120 (2023).

36 Eigenbrood, J. et al. Spatial Profiling Identifies Regionally Distinct Microenvironments and Targetable Immunosuppressive Mechanisms in Pediatric Osteosarcoma Pulmonary Metastases. Cancer Res 85, 2320–2337 (2025).

37 Alshebremi, M. et al. Functional tumor cell-intrinsic STING, not host STING, drives local and systemic antitumor immunity and therapy efficacy following cryoablation. J Immunother Cancer 11 (2023).

38 Sen, T. et al. Targeting DNA Damage Response Promotes Antitumor Immunity through STING-Mediated T-cell Activation in Small Cell Lung Cancer. Cancer Discov 9, 646–661 (2019).

39 Ohkuri, T. et al. Intratumoral administration of cGAMP transiently accumulates potent macrophages for anti-tumor immunity at a mouse tumor site. Cancer Immunol Immunother 66, 705–716 (2017).

40. Jneid, B., et al. Selective STING stimulation in dendritic cells primes antitumor T cell responses. Sci Immunol 8, eabn6612 (2023).

41 Corrales, L. et al. Direct Activation of STING in the Tumor Microenvironment Leads to Potent and Systemic Tumor Regression and Immunity. Cell reports 11, 1018–1030 (2015).

42 Sayles, L. C. et al. Genome-Informed Targeted Therapy for Osteosarcoma. Cancer Discov 9, 46–63 (2019).

43 Toda, G. et al. Preparation and culture of bone marrow-derived macrophages from mice for functional analysis. STAR Protoc 2, 100246 (2021).

44 Takeshita, S. et al. Identification and characterization of the new osteoclast progenitor with macrophage phenotypes being able to differentiate into mature osteoclasts. J Bone Miner Res 15, 1477–1488 (2000).

45 Bolger, A. M. et al. Trimmomatic: a flexible trimmer for Illumina sequence data. Bioinformatics 30, 2114–2120 (2014).

46 Dobin, A. et al. STAR: ultrafast universal RNA-seq aligner. Bioinformatics 29, 15–21 (2013).

47 Robinson, M. D. et al. edgeR: a Bioconductor package for differential expression analysis of digital gene expression data. Bioinformatics 26, 139–140 (2010).

48 Law, C. W. et al. voom: Precision weights unlock linear model analysis tools for RNA-seq read counts. Genome Biol 15, R29 (2014).

49 Newman, A. M. et al. Determining cell type abundance and expression from bulk tissues with digital cytometry. Nat Biotechnol 37, 773–782 (2019).

50 Niu, B. et al. MSIsensor: microsatellite instability detection using paired tumor-normal sequence data. Bioinformatics 30, 1015–1016 (2014).

51 Yoshihara, K. et al. Inferring tumour purity and stromal and immune cell admixture from expression data. Nat Commun 4, 2612 (2013).

52 Newman, A. M. et al. Robust enumeration of cell subsets from tissue expression profiles. Nat Methods 12, 453–457 (2015).

53 Vivian, J. et al. Toil enables reproducible, open source, big biomedical data analyses. Nat Biotechnol 35, 314–316 (2017).

