## Supplementary material for "STING activation reshapes the tumor microenvironment leading to tumor regression in osteosarcoma": All Supplemental Information

### Fig. S1

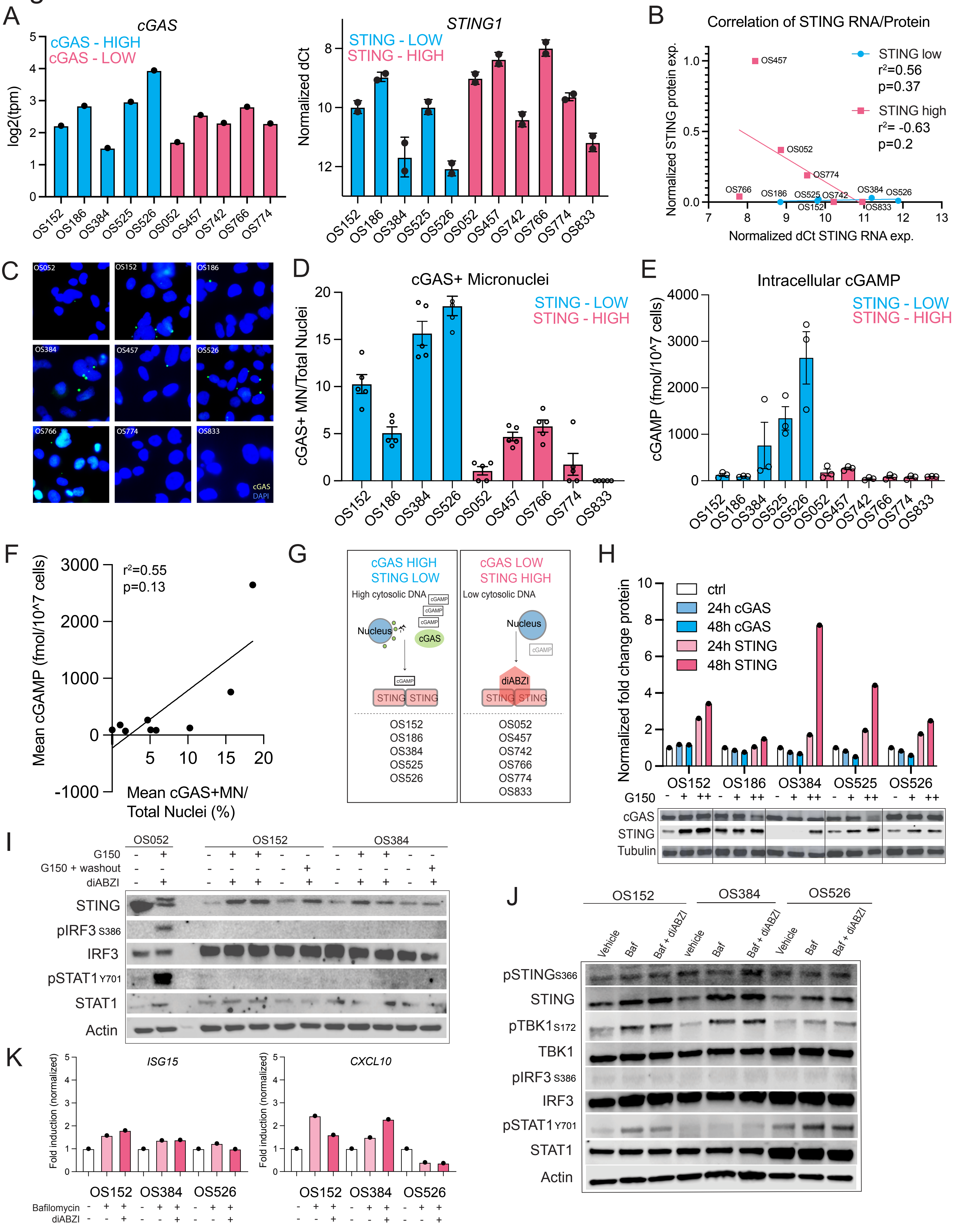
A
cGAS
cGAS - HIGH
cGAS - LOW
log2(tpm)
OS152 OS186 OS384 OS525 OS526 OS052 OS457 OS742 OS766 OS774
STING1
STING - LOW
STING - HIGH
Normalized dCt
OS152 OS186 OS384 OS525 OS526 OS052 OS457 OS742 OS766 OS774 OS833
B
Correlation of STING RNA/Protein
STING low
r²=0.56
p=0.37
STING high
r²= -0.63
p=0.2
Normalized STING protein exp.
Normalized dCt STING RNA exp.
C
cGAS
DAPI
D
cGAS+ Micronuclei
STING - LOW
STING - HIGH
cGAS+ MN/Total Nuclei
E
Intracellular cGAMP
STING - LOW
STING - HIGH
cGAMP (fmol/10^7 cells)
F
r²=0.55
p=0.13
Mean cGAMP (fmol/10^7 cells)
Mean cGAS+MN/
Total Nuclei (%)
G
cGAS HIGH
STING LOW
High cytosolic DNA
Nucleus
cGAS
cGAMP
STING STING
OS152
OS186
OS384
OS525
OS526
cGAS LOW
STING HIGH
Low cytosolic DNA
Nucleus
cGAMP
diABZI
STING STING
OS052
OS457
OS742
OS766
OS774
OS833
H
ctrl
24h cGAS
48h cGAS
24h STING
48h STING
Normalized fold change protein
OS152 OS186 OS384 OS525 OS526
G150 - + ++
cGAS
STING
Tubulin
I
OS052 OS152 OS384
G150
G150 + washout
diABZI
STING
pIRF3 S386
IRF3
pSTAT1 Y701
STAT1
Actin
J
OS152 OS384 OS526
Vehicle
Baf
Baf + diABZI
pSTING S366
STING
pTBK1 S172
TBK1
pIRF3 S386
IRF3
pSTAT1 Y701
STAT1
Actin
K
ISG15
CXCL10
Fold induction (normalized)
OS152 OS384 OS526
Bafilomycin
diABZI

**Figure S1. S1A.** CGAS transcript analysis by bulk RNAseq and STING1 transcript analysis by qPCR in osteosarcoma PDX cell lines. **S1B.** Spearman correlation of STING1 RNA expression by qPCR (Fig. S1A) and protein expression quantified (Fig. 1B) for STING-low/high groups separately. **S1C-D.** Micronuclei (MN) are identified in all osteosarcoma cell lines but vary in overall burden and cGAS recruitment to MN. (C) Representative IF images demonstrating MN (DAPI) and cGAS (FITC) across osteosarcoma PDX cell line panel. (D) Manual quantification of percentage of MN with co-positivity for cGAS of total nuclei (mean ± SEM; n=5 FOV/cell line). **S1E.** Intracellular cGAMP as measured by ELISA across panel of osteosarcoma PDX cell lines (mean ± SEM; n=3 per cell line across 3 independent experiments). S1F. Spearman correlation of cGAS+ MN burden by IF (Fig. S1D) and intracellular cGAMP production (Fig. S1E) in osteosarcoma PDX cell lines (n=9). **S1G.** Schema classifying osteosarcoma PDX cell lines based on characterization of cGAS/STING protein levels and response to STING agonist treatment. **S1H.** Treatment of STING-low PDX cell lines with cGAS inhibitor increases total STING protein levels. **S1I.** Immunoblot analysis for STING activation after combination treatment with cGAS inhibitor (± washout) and STING agonist (diABZI). **S1J-K.** Treatment of STING-low PDX cell lines with bafilomycin and diABZI for 6h and profiling of STING activation by immunoblotting (J) and qPCR (K). Cell lines are color coded according to STING protein in 1B as STING-high (pink) or STING-low (blue).

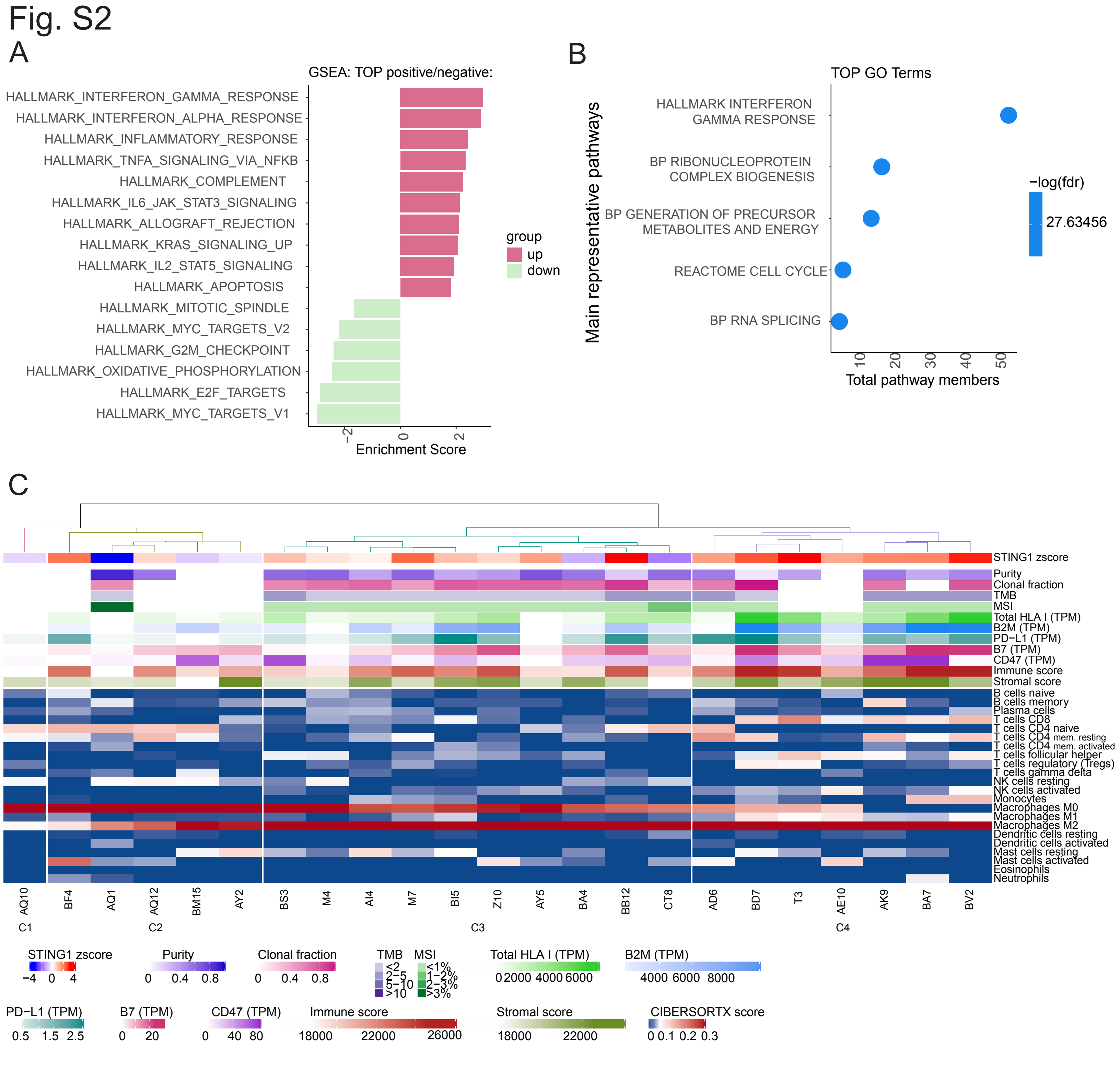


**Figure S2. S2A-B.** Pathway analysis of STING activation signature (Fig. 2B). **S2C.** Immuno-profiling was performed on osteosarcoma diagnostic biopsy tissue samples (n=23) using CIBERSORTX and annotated with genomic features from RNAseq and WGS data. Missing genomic annotations indicate samples where WGS was unavailable. Based on the immune cell type scores, samples were clustered into 4 groups using k-means clustering.

Fig. S3

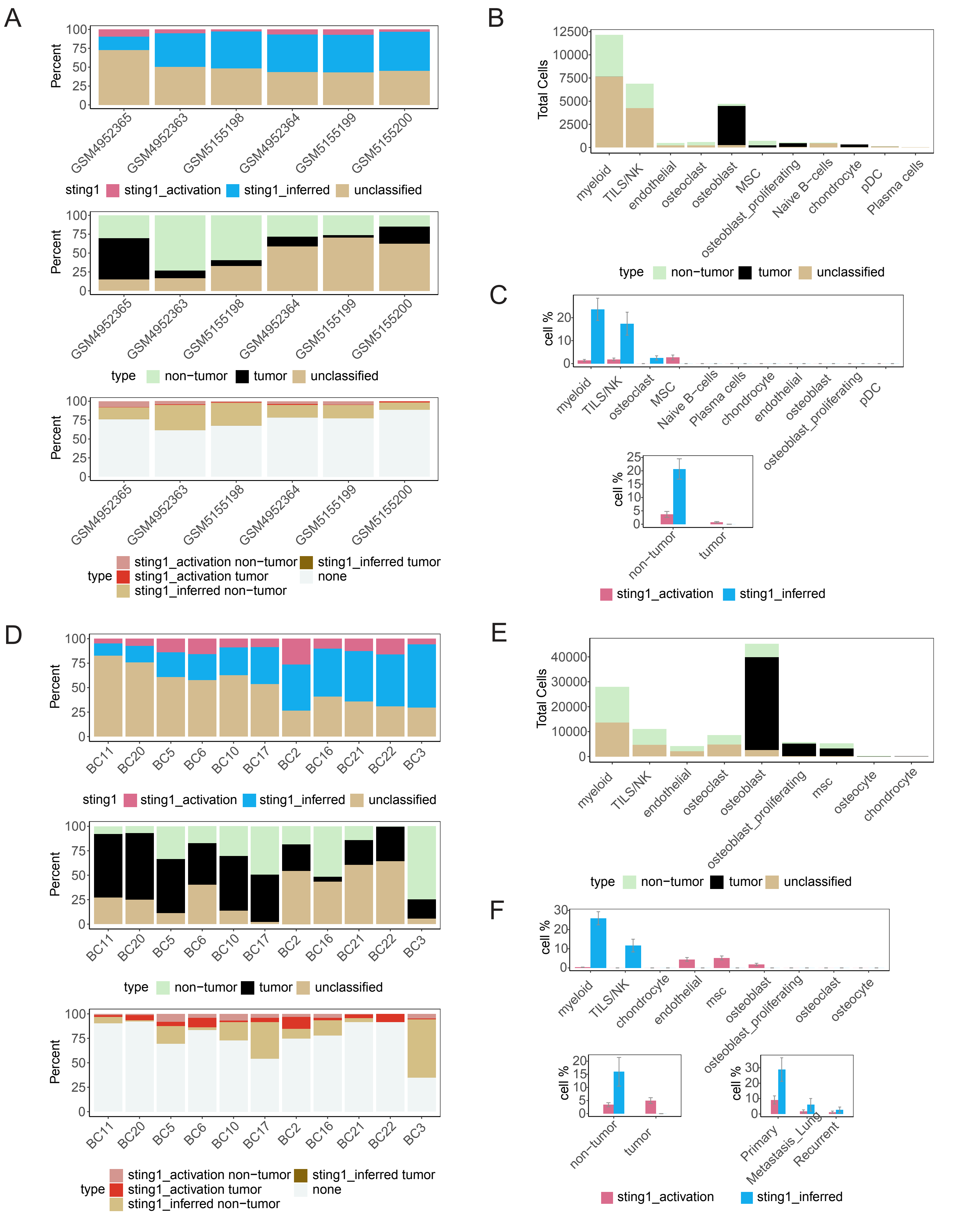

A
Percent
100
75
50
25
0
GSM4952365
GSM4952363
GSM5155198
GSM4952364
GSM5155199
GSM5155200
sting1
sting1_activation
sting1_inferred
unclassified
type
non−tumor
tumor
unclassified
sting1_activation non−tumor
sting1_activation tumor
sting1_inferred non−tumor
sting1_inferred tumor
none
B
Total Cells
12500
10000
7500
5000
2500
0
myeloid
TILS/NK
endothelial
osteoclast
osteoblast
MSC
osteoblast_proliferating
Naive B−cells
chondrocyte
pDC
Plasma cells
type
non−tumor
tumor
unclassified
C
cell %
20
10
0
myeloid
TILS/NK
osteoclast
MSC
Naive B−cells
Plasma cells
chondrocyte
endothelial
osteoblast
osteoblast_proliferating
pDC
cell %
25
20
15
10
5
0
non−tumor
tumor
sting1_activation
sting1_inferred
D
Percent
100
75
50
25
0
BC11
BC20
BC5
BC6
BC10
BC17
BC2
BC16
BC21
BC22
BC3
sting1
sting1_activation
sting1_inferred
unclassified
type
non−tumor
tumor
unclassified
type
sting1_activation non−tumor
sting1_activation tumor
sting1_inferred non−tumor
sting1_inferred tumor
none
E
Total Cells
40000
30000
20000
10000
0
myeloid
TILS/NK
endothelial
osteoclast
osteoblast
osteoblast_proliferating
msc
osteocyte
chondrocyte
type
non−tumor
tumor
unclassified
F
cell %
30
20
10
0
myeloid
TILS/NK
chondrocyte
endothelial
msc
osteoblast
osteoblast_proliferating
osteoclast
osteocyte
cell %
20
15
10
5
0
non−tumor
tumor
cell %
30
20
10
0
Primary
Metastasis_Lung
Recurrent
sting1_activation
sting1_inferred

**Figure S3.** Supporting bar plots for scRNAseq analysis in Fig. 3. Dataset of 6 treatment-naïve primary osteosarcoma samples is included in S3A-C (25) and dataset of 11 treated osteosarcoma samples (9 primary and 2 metastatic) is included in S3D-F (24). **S3A, D**. Stackplots of STING activation and STING inferred enrichment by sample (top), tumor vs. normal by sample (middle), and sub-setting of STING signatures in tumor vs. normal by sample (bottom). **S3B-C, E-F**. Bar plots indicating percentage of cells in indicated categories by cell type.

### Fig. S4

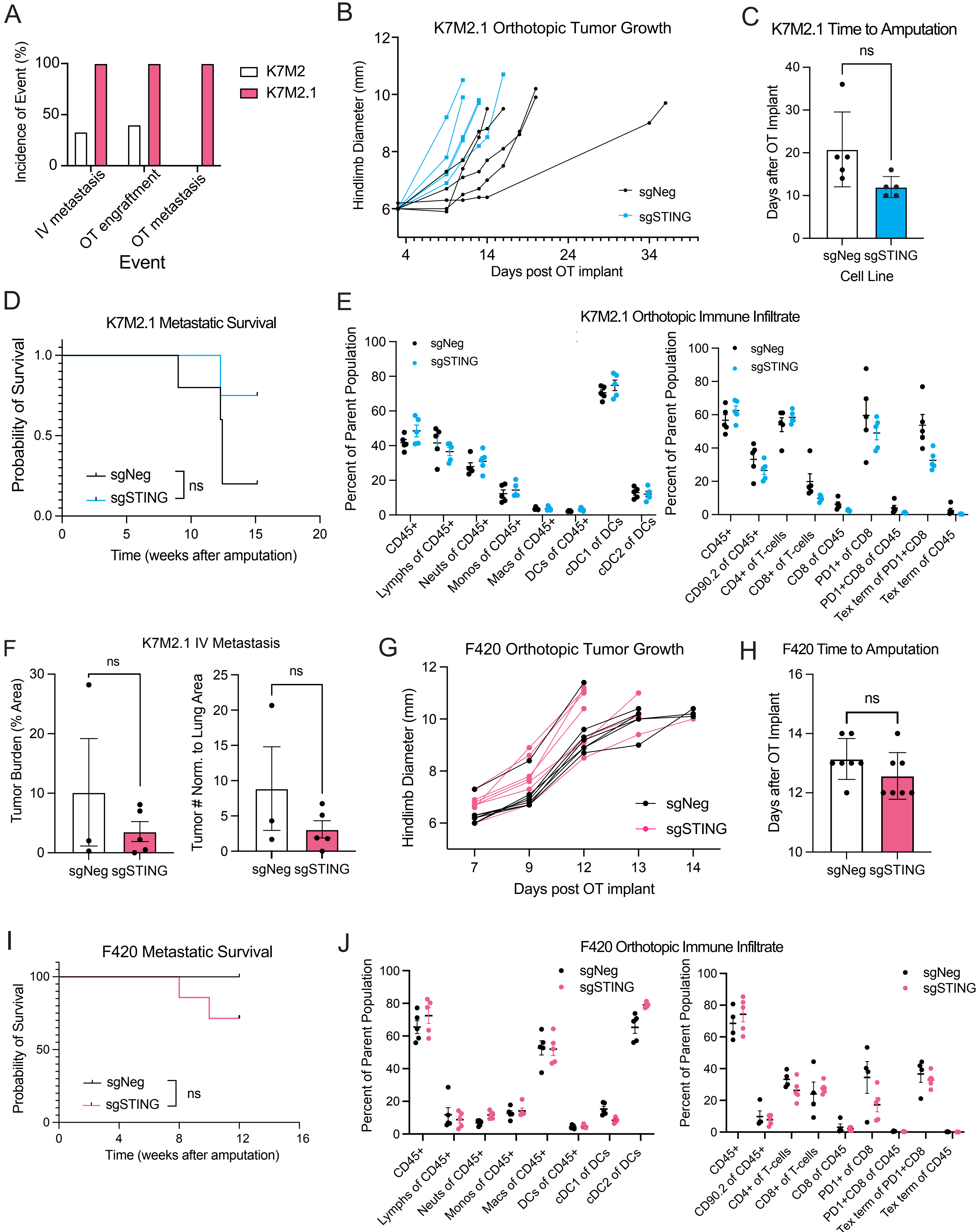
A
Incidence of Event (%)
100
50
0
K7M2
K7M2.1
IV metastasis
OT engraftment
OT metastasis
Event
B
K7M2.1 Orthotopic Tumor Growth
Hindlimb Diameter (mm)
12
10
8
6
4
14
24
34
Days post OT implant
sgNeg
sgSTING
C
K7M2.1 Time to Amputation
ns
Days after OT Implant
40
30
20
10
0
sgNeg sgSTING
Cell Line
D
K7M2.1 Metastatic Survival
Probability of Survival
1.0
0.5
0.0
0
5
10
15
20
Time (weeks after amputation)
sgNeg
sgSTING
ns
E
K7M2.1 Orthotopic Immune Infiltrate
Percent of Parent Population
100
80
60
40
20
0
sgNeg
sgSTING
CD45+
Lymphs of CD45+
Neuts of CD45+
Monos of CD45+
Macs of CD45+
DCs of CD45+
cDC1 of DCs
cDC2 of DCs
CD45+
CD90.2 of CD45+
CD4+ of T-cells
CD8+ of T-cells
CD8 of CD45
PD1+ of CD8
PD1+CD8 of CD45
Tex term of PD1+CD8
Tex term of CD45
F
K7M2.1 IV Metastasis
Tumor Burden (% Area)
30
20
10
0
ns
Tumor # Norm. to Lung Area
25
20
15
10
5
0
sgNeg sgSTING
G
F420 Orthotopic Tumor Growth
Hindlimb Diameter (mm)
12
10
8
6
7
9
12
13
14
Days post OT implant
sgNeg
sgSTING
H
F420 Time to Amputation
ns
Days after OT Implant
16
14
12
10
sgNeg sgSTING
I
F420 Metastatic Survival
Probability of Survival
100
50
0
0
4
8
12
16
Time (weeks after amputation)
sgNeg
sgSTING
ns
J
F420 Orthotopic Immune Infiltrate
Percent of Parent Population
100
80
60
40
20
0
sgNeg
sgSTING
CD45+
Lymphs of CD45+
Neuts of CD45+
Monos of CD45+
Macs of CD45+
DCs of CD45+
cDC1 of DCs
cDC2 of DCs
CD45+
CD90.2 of CD45+
CD4+ of T-cells
CD8+ of T-cells
CD8 of CD45
PD1+ of CD8
PD1+CD8 of CD45
Tex term of PD1+CD8
Tex term of CD45

**Figure S4. S4A.** Incidence of tumor engraftment and metastasis comparing parental K7M2 to K7M2.1, which was derived from a K7M2 OT tumor. IV studies with K7M2 (n=6 in 2 experiments) reached a 4-6 week endpoint; K7M2.1 mice (n=4 in 1 experiment) reached endpoint at 3 weeks. OT studies were monitored for engraftment for 10-18 weeks, with a metastatic endpoint of 6-8 weeks post-amputation; K7M2 (n=5 in 2 experiments), K7M2.1 (n=3 in 1 experiment). **S4B-C.** Primary tumor growth as measured by time to amputation for K7M2.1 sgNeg (n=5) vs. sgSTING (n=5); **S4D.** Survival after amputation for K7M2.1 sgNeg (n=5) vs. sgSTING (n=4); **S4E.** Immune infiltration by flow cytometry (left, myeloid panel; right, lymphoid panel) in OT osteosarcoma tumors, K7M2.1 sgNeg (n=5) and sgSTING (n=5). **S4F.** IV injection of K7M2.1 sgNeg (n=3) and sgSTING (n=5) cells in BALB/c mice with histopathologic quantification of tumor burden at endpoint. **S4G-I.** Murine osteosarcoma cell line F420 sgNeg (n=7) and sgSTING (n=7) primary tumor growth as measured by time to amputation and metastatic survival. **S4J.** Immune infiltration by flow cytometry (left, myeloid panel; right, lymphoid panel) in F420 sgNeg (n=5 for myeloid panel; n=4 for lymphoid panel) and sgSTING (n=5) tumors. Data are shown as mean (A), mean ± SD (C, H) and mean ± SEM (E, F, J); significant differences were not observed; unpaired t test (C, F, H), unpaired t test followed by two-stage step-up multiple comparisons test performed separately for myeloid and lymphoid panels (E, J) and log-rank (Mantel-Cox) test (D and I).

### Fig. S5

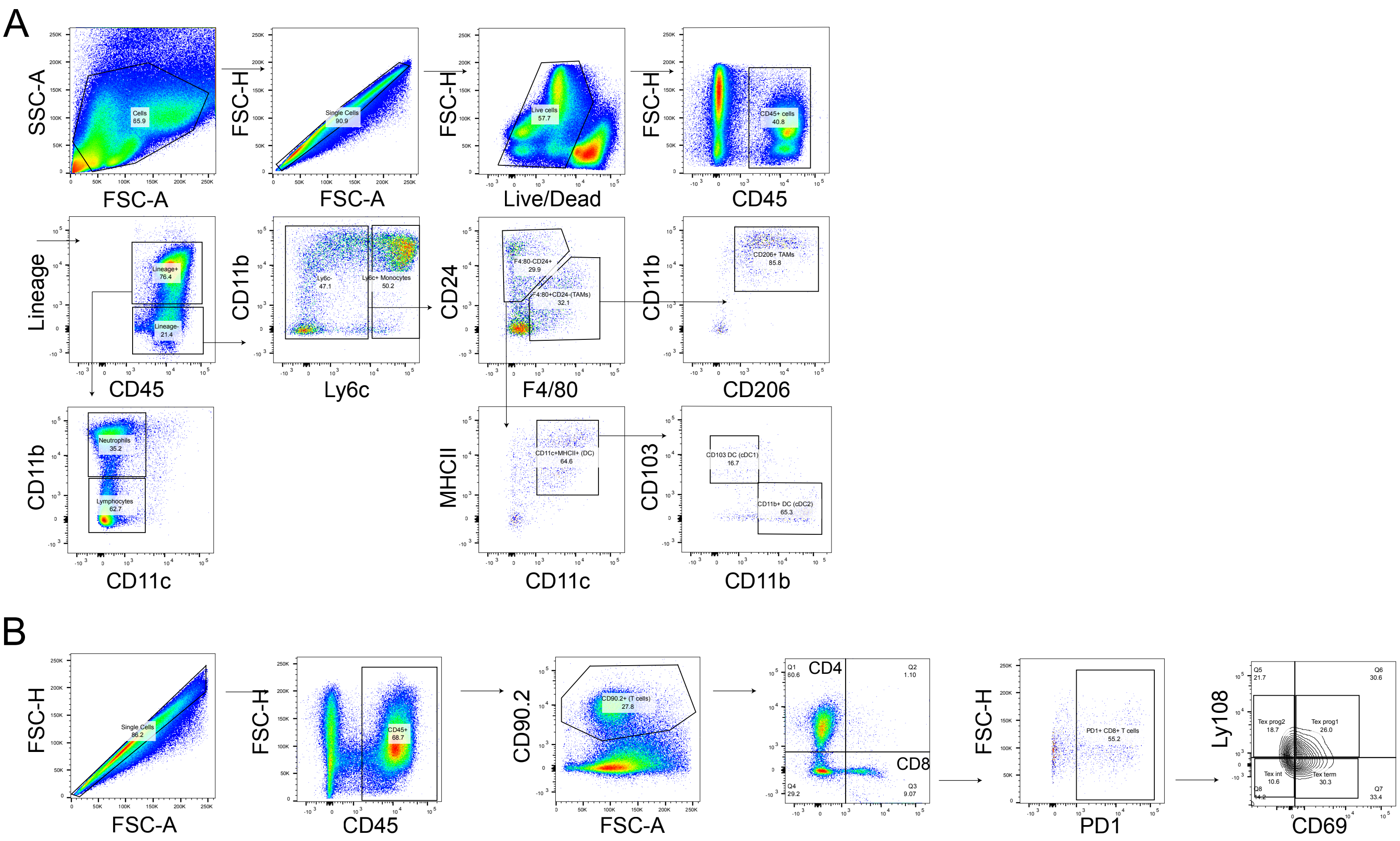


**Figure S5. Representative gating strategy of murine osteosarcoma tumors.** Fresh or viably frozen OT tumors were digested and partitioned for myeloid (S4A) and T lymphoid (S4B) panels. **S4A.** Immunophenotyping for myeloid markers within lineage (CD90.2, CD45R/B220, Ly-6G, NK-1.1, Siglec-F) negative population, including CD11b, Ly6c, CD24, F4/80, CD206, MHCII, CD11c, and CD103. For flow cytometry of BALB/c-derived tumors, NK cells were not included in lineage-positive population and where specified a distinct NK marker (CD49b) was utilized to characterize NK cell changes after gating out lineage-positive cells. **S4B.** Immunophenotyping for T lymphoid markers within CD90.2 positive gate, including CD4, CD8, PD1, Ly108, and CD69.

**Supplemental Table 1**

| gene | logFC | P.Value | adj.P.Val | treated.ave | control.ave | class |
|---|---|---|---|---|---|---|
| AC005532.1 | 1.8 | 0.0041 | 0.0129 | 2.5115 | 1.8611 | up |
| AC023157.1 | 2.27 | 0 | 0 | 1.7921 | 0.6277 | up |
| AGRN | 1.54 | 0 | 0 | 7.3401 | 5.6254 | up |
| AL365181.3 | 1.56 | 0 | 0.0001 | 2.5547 | 1.6182 | up |
| APOL6 | 2.1 | 0.0168 | 0.0413 | 5.5551 | 2.8843 | up |
| BST2 | 2.65 | 0 | 0.0002 | 7.375 | 2.8984 | up |
| CCL5 | 4.88 | 0 | 0 | 4.5673 | 0.7645 | up |
| CMPK2 | 5.45 | 0 | 0 | 4.8337 | 1.0028 | up |
| CXCL10 | 3.62 | 0 | 0 | 1.0705 | 0.096 | up |
| CYP24A1 | 2.09 | 0 | 0.0002 | 2.3285 | 1.4181 | up |
| DDX58 | 2.78 | 0 | 0 | 6.631 | 3.9317 | up |
| DDX60 | 3.14 | 0 | 0 | 6.6026 | 2.963 | up |
| DDX60L | 2.89 | 0 | 0 | 5.9101 | 2.669 | up |
| DHRS2 | 1.82 | 0 | 0 | 4.3594 | 3.1315 | up |
| DHX58 | 3.43 | 0 | 0 | 3.5022 | 1.0294 | up |
| DSP | 1.55 | 0.0054 | 0.0161 | 2.8646 | 1.5131 | up |
| DTX3L | 2.63 | 0 | 0 | 7.4483 | 4.7846 | up |
| EIF2AK2 | 1.68 | 0 | 0 | 8.6115 | 6.9054 | up |
| EPSTI1 | 2.69 | 0 | 0 | 4.1151 | 1.7065 | up |
| FBXO6 | 1.68 | 0 | 0 | 1.9841 | 0.9258 | up |
| FER1L6 | 1.63 | 0.0001 | 0.0007 | 2.8746 | 1.9092 | up |
| FLT1 | 1.7 | 0.0014 | 0.0052 | 2.266 | 1.4435 | up |
| FST | 1.6 | 0 | 0 | 3.9244 | 2.8973 | up |
| GBP1 | 2.42 | 0 | 0 | 6.2408 | 3.6002 | up |
| HCP5 | 1.7 | 0.0002 | 0.0012 | 3.2594 | 1.7185 | up |
| HELZ2 | 2.85 | 0 | 0 | 7.122 | 4.2566 | up |
| HERC5 | 2.41 | 0 | 0 | 5.8941 | 3.0274 | up |
| HERC6 | 4.38 | 0 | 0 | 6.8751 | 2.727 | up |
| HES4 | 2.65 | 0 | 0 | 2.675 | 0.8795 | up |
| HLA-B | 1.83 | 0 | 0 | 8.5229 | 6.6319 | up |
| HLA-C | 1.52 | 0 | 0 | 8.7682 | 7.0792 | up |
| HLA-F | 2.2 | 0 | 0 | 4.6248 | 2.5354 | up |
| IFI27 | 4.51 | 0 | 0 | 5.895 | 2.0769 | up |
| IFI35 | 3.03 | 0.0001 | 0.0005 | 4.6927 | 1.7215 | up |
| IFI44 | 3.45 | 0 | 0 | 7.8014 | 3.76 | up |
| IFI44L | 4.84 | 0 | 0 | 7.7309 | 2.1554 | up |
| IFI6 | 4.83 | 0 | 0 | 10.0725 | 5.1057 | up |
| IFIH1 | 3.64 | 0 | 0 | 6.0946 | 2.7333 | up |
| IFIT1 | 3.53 | 0 | 0 | 7.8836 | 4.354 | up |
| IFIT2 | 2.8 | 0 | 0 | 6.9978 | 4.1439 | up |
| IFIT3 | 3.39 | 0 | 0 | 8.0031 | 4.4641 | up |
| IFITM1 | 4.22 | 0 | 0 | 6.0601 | 1.8895 | up |
| IFITM3 | 2.56 | 0 | 0 | 9.4327 | 6.6651 | up |
| IL7R | 1.52 | 0 | 0.0001 | 5.7109 | 3.9982 | up |
| IRF1-AS1 | 1.54 | 0 | 0 | 2.4354 | 1.385 | up |
| IRF7 | 3.49 | 0 | 0 | 5.6105 | 2.3826 | up |

| | | | | | | |
|---|---|---|---|---|---|---|
| IRF9 | 1.66 | 0 | 0 | 2.2852 | 1.144 | up |
| ISG15 | 5.22 | 0 | 0 | 9.3388 | 4.16 | up |
| ISG20 | 2.05 | 0 | 0 | 3.4803 | 1.4811 | up |
| LGALS3BP | 2.18 | 0 | 0.0002 | 7.9558 | 5.2786 | up |
| LIF | 1.64 | 0 | 0 | 2.8346 | 1.8284 | up |
| LY6E | 1.88 | 0 | 0 | 8.1771 | 5.9234 | up |
| MX1 | 5.41 | 0 | 0 | 8.8948 | 3.0661 | up |
| MX2 | 2.96 | 0 | 0.0002 | 5.2151 | 1.8343 | up |
| NMI | 1.57 | 0.005 | 0.0153 | 4.0824 | 2.4093 | up |
| NT5E | 1.72 | 0 | 0 | 5.3838 | 3.7499 | up |
| OAS1 | 5.64 | 0 | 0 | 6.952 | 1.8825 | up |
| OAS2 | 5.28 | 0 | 0 | 7.7076 | 2.2996 | up |
| OAS3 | 4.11 | 0 | 0 | 9.4648 | 5.3464 | up |
| OASL | 3.07 | 0 | 0 | 6.2152 | 2.6692 | up |
| PARP10 | 2.66 | 0.0001 | 0.0004 | 5.2778 | 2.1954 | up |
| PARP12 | 2.86 | 0 | 0 | 6.0047 | 3.4629 | up |
| PARP14 | 2.73 | 0 | 0 | 7.6444 | 4.9747 | up |
| PARP9 | 3.02 | 0 | 0 | 7.2411 | 4.2058 | up |
| PLAAT4 | 3 | 0 | 0 | 3.8639 | 1.4662 | up |
| PLEKHA4 | 2.04 | 0 | 0.0001 | 4.345 | 2.6714 | up |
| PLSCR1 | 2.26 | 0 | 0 | 7.0404 | 4.7873 | up |
| PSMB9 | 2.17 | 0.0001 | 0.0005 | 4.2214 | 2.183 | up |
| PTGS2 | 1.79 | 0 | 0 | 2.8298 | 1.9583 | up |
| REC8 | 1.89 | 0 | 0 | 3.1391 | 1.5892 | up |
| RSAD2 | 6.57 | 0 | 0 | 5.3352 | 1.1969 | up |
| RXRG | 3.81 | 0 | 0 | 3.0612 | 0.631 | up |
| SAMD9 | 3.16 | 0 | 0 | 7.515 | 4.5565 | up |
| SAMD9L | 3.32 | 0 | 0 | 6.3787 | 3.1602 | up |
| SAMHD1 | 2.22 | 0 | 0 | 7.5282 | 5.3832 | up |
| SERPING1 | 2.83 | 0.0005 | 0.0022 | 3.1449 | 1.5816 | up |
| SHFL | 2.48 | 0 | 0 | 6.0284 | 3.8288 | up |
| SP100 | 1.69 | 0 | 0 | 6.586 | 5.0624 | up |
| SP110 | 2.61 | 0 | 0 | 3.4996 | 1.6129 | up |
| STAT1 | 2.14 | 0 | 0 | 9.0841 | 6.9764 | up |
| STAT2 | 1.67 | 0 | 0 | 7.2945 | 5.752 | up |
| TAP1 | 1.64 | 0 | 0 | 6.9761 | 5.3142 | up |
| TEX41 | 1.63 | 0 | 0 | 2.9866 | 2.2628 | up |
| TLR3 | 1.61 | 0 | 0 | 3.6953 | 2.1424 | up |
| TM7SF2 | 1.81 | 0 | 0 | 2.3655 | 1.6029 | up |
| TMEM140 | 2.13 | 0 | 0 | 2.5332 | 0.9229 | up |
| TRANK1 | 1.86 | 0 | 0.0001 | 3.8698 | 1.8848 | up |
| TRIM21 | 1.62 | 0 | 0 | 4.8218 | 3.4085 | up |
| TRIM22 | 4 | 0 | 0 | 3.8086 | 0.8635 | up |
| TYMP | 2.02 | 0 | 0 | 2.1525 | 1.0419 | up |
| UBA7 | 2.54 | 0 | 0 | 4.354 | 1.7071 | up |

| | | | | | | |
|---|---|---|---|---|---|---|
| UBE2L6 | 2.85 | 0 | 0 | 5.963 | 3.2269 | up |
| USP18 | 3.04 | 0 | 0 | 5.5751 | 2.9724 | up |
| XAF1 | 4.91 | 0 | 0 | 4.5755 | 1.252 | up |
| TNFRSF10D | -1.61 | 0 | 0.0001 | 2.0456 | 3.1073 | down |

**Supplemental Table 2**

| gene | logFC | AveExpr | t | P.Value | adj.P.Val |
|---|---|---|---|---|---|
| LCP1 | 2.74 | 6.73 | 5.86 | 0 | 0.0157 |
| C1QB | 4.03 | 6.61 | 5.69 | 0 | 0.0157 |
| C1QA | 3.52 | 6.61 | 5.56 | 0 | 0.0157 |
| CABP1 | -3.66 | 0.33 | -5.48 | 0 | 0.0157 |
| SIGLEC1 | 3.14 | 4.38 | 5.35 | 0 | 0.0157 |
| PYGL | -2.82 | 6.26 | -5.34 | 0 | 0.0157 |
| C1QC | 3.99 | 6.42 | 5.33 | 0 | 0.0157 |
| SLC15A3 | 2.64 | 4.13 | 5.14 | 0 | 0.0157 |
| ENC1 | 2.37 | 4.52 | 5.13 | 0 | 0.0157 |
| APOE | 3.4 | 8 | 5.13 | 0 | 0.0157 |
| CD7 | 5.32 | 0.01 | 5.13 | 0 | 0.0157 |
| LGALS9 | 2.65 | 5.23 | 5.11 | 0 | 0.0157 |
| SEPTIN6 | 1.77 | 4.71 | 5.1 | 0 | 0.0157 |
| GZMK | 5.52 | 0.52 | 5.08 | 0 | 0.0157 |
| HLA-DMA | 2.8 | 5.4 | 5.02 | 0 | 0.0157 |
| ARG2 | -3.09 | 2.43 | -5.02 | 0 | 0.0157 |
| ADA2 | 3.31 | 4.01 | 5.01 | 0 | 0.0157 |
| CD74 | 3.39 | 9.62 | 5.01 | 0 | 0.0157 |
| IL18BP | 2.28 | 3.47 | 4.94 | 0 | 0.0168 |
| GBP4 | 4.01 | 3.98 | 4.94 | 0 | 0.0168 |
| HCLS1 | 2.24 | 5.7 | 4.92 | 0 | 0.0168 |
| CYTH4 | 2.72 | 5.08 | 4.89 | 0 | 0.0168 |
| CD2 | 6.08 | 1.14 | 4.89 | 0 | 0.0168 |
| SRPK3 | -5.06 | 0.75 | -4.86 | 0 | 0.0168 |
| SERPINB9 | 3.03 | 2.83 | 4.85 | 0 | 0.0168 |
| PSME2 | 1.63 | 6.23 | 4.85 | 0 | 0.0168 |
| IFIT5 | 1.46 | 3.93 | 4.85 | 0 | 0.0168 |
| PRF1 | 3.91 | 1.22 | 4.79 | 0 | 0.019 |
| CD48 | 3.37 | 2.84 | 4.76 | 0 | 0.0195 |
| CAMSAP1 | -1.19 | 5.16 | -4.76 | 0 | 0.0195 |
| MS4A6A | 2.95 | 5.35 | 4.71 | 0 | 0.0203 |
| TRBC1 | 5.61 | 0.4 | 4.71 | 0 | 0.0203 |
| CD5 | 4.81 | 0.09 | 4.71 | 0 | 0.0203 |
| SLA2 | 4.49 | -0.09 | 4.7 | 0.0001 | 0.0203 |
| SIPA1 | 1.62 | 5.29 | 4.68 | 0.0001 | 0.0211 |
| B2M | 2.08 | 10.55 | 4.66 | 0.0001 | 0.0215 |
| GIMAP4 | 2.6 | 4.49 | 4.63 | 0.0001 | 0.0222 |
| MACROH2A2 | 2.09 | 3.95 | 4.63 | 0.0001 | 0.0222 |
| SLA | 3.05 | 3.5 | 4.61 | 0.0001 | 0.0222 |
| LILRB4 | 3.17 | 4.73 | 4.61 | 0.0001 | 0.0222 |
| HLA-DRA | 3.1 | 8.45 | 4.6 | 0.0001 | 0.0222 |
| SLAMF6 | 4.22 | 0.7 | 4.57 | 0.0001 | 0.0235 |
| HLA-DPB1 | 3.06 | 6.93 | 4.57 | 0.0001 | 0.0235 |

| | | | | | |
|---|---|---|---|---|---|
| PSME1 | 1.29 | 6.5 | 4.55 | 0.0001 | 0.0241 |
| PLCB2 | 3.03 | 4.91 | 4.54 | 0.0001 | 0.0241 |
| DENND1C | 3.07 | 2.66 | 4.51 | 0.0001 | 0.0245 |
| CD52 | 4.31 | 2.29 | 4.51 | 0.0001 | 0.0245 |
| TP63 | -3.67 | 2.12 | -4.51 | 0.0001 | 0.0245 |
| NCF4 | 2.6 | 3.83 | 4.5 | 0.0001 | 0.0245 |
| CD4 | 2.97 | 6.12 | 4.5 | 0.0001 | 0.0246 |
| IL2RG | 3.99 | 3.04 | 4.49 | 0.0001 | 0.0249 |
| MILR1 | 2.41 | 2.89 | 4.47 | 0.0001 | 0.0253 |
| AACS | -1.61 | 4.15 | -4.46 | 0.0001 | 0.0255 |
| TRBC2 | 5.16 | 1.55 | 4.46 | 0.0001 | 0.0255 |
| RASAL3 | 2.85 | 2.63 | 4.44 | 0.0001 | 0.0259 |
| EVI2B | 3.02 | 4.23 | 4.44 | 0.0001 | 0.0259 |
| TMEM176B | 3.76 | 4.82 | 4.43 | 0.0001 | 0.0262 |
| ARHGAP4 | 3.03 | 4.55 | 4.42 | 0.0001 | 0.0262 |
| CD6 | 5.11 | 0.54 | 4.42 | 0.0001 | 0.0262 |
| ARHGAP9 | 2.53 | 3.45 | 4.39 | 0.0001 | 0.027 |
| RTP4 | 4.25 | 0.24 | 4.39 | 0.0001 | 0.027 |
| CLEC4E | 4.32 | -0.08 | 4.39 | 0.0001 | 0.027 |
| ADAMTS7 | 1.89 | 5.25 | 4.36 | 0.0001 | 0.028 |
| CA5B | -2.28 | 3.22 | -4.36 | 0.0001 | 0.028 |
| FCER1G | 3.04 | 5.97 | 4.36 | 0.0001 | 0.028 |
| LTB | 5.14 | 0.33 | 4.35 | 0.0001 | 0.028 |
| SASH3 | 3.09 | 2.65 | 4.35 | 0.0001 | 0.0281 |
| LST1 | 3.24 | 3.09 | 4.34 | 0.0001 | 0.0285 |
| TMEM150B | 3.95 | -0.04 | 4.33 | 0.0001 | 0.029 |
| BTN3A2 | 1.83 | 5.65 | 4.3 | 0.0002 | 0.0295 |
| RASSF4 | 2.69 | 4.88 | 4.3 | 0.0002 | 0.0295 |
| PTPN7 | 4.08 | 2.45 | 4.3 | 0.0002 | 0.0295 |
| FHOD3 | -1.89 | 5.27 | -4.3 | 0.0002 | 0.0295 |
| LPXN | 2.23 | 3.37 | 4.27 | 0.0002 | 0.0315 |
| CYTH1 | 1.24 | 5.33 | 4.26 | 0.0002 | 0.0324 |
| MYH3 | -3.87 | 2.76 | -4.24 | 0.0002 | 0.0326 |
| HCK | 2.89 | 4.13 | 4.23 | 0.0002 | 0.0326 |
| APOC1 | 3.08 | 3.55 | 4.23 | 0.0002 | 0.0326 |
| SECTM1 | 3.78 | 2.23 | 4.23 | 0.0002 | 0.0326 |
| KLF5 | -2.48 | 3.12 | -4.23 | 0.0002 | 0.0326 |
| NT5C2 | 1.09 | 5.6 | 4.22 | 0.0002 | 0.0326 |
| SCML2 | -2.44 | 2.17 | -4.22 | 0.0002 | 0.0326 |
| ARHGDIB | 2.51 | 6.51 | 4.21 | 0.0002 | 0.0333 |
| CCNJL | 2.5 | 1.83 | 4.2 | 0.0002 | 0.0336 |
| FERMT3 | 2.4 | 5.02 | 4.19 | 0.0002 | 0.0344 |
| PRRC2B | -1.28 | 7.77 | -4.18 | 0.0002 | 0.0346 |
| NABP1 | 1.49 | 4.23 | 4.18 | 0.0002 | 0.0346 |
| CXCL11 | 5.73 | 0.09 | 4.18 | 0.0002 | 0.0346 |

| | | | | | |
|---|---|---|---|---|---|
| FAM160A1 | -3.22 | 1 | -4.16 | 0.0002 | 0.0357 |
| SLC38A5 | -2.64 | 4.26 | -4.15 | 0.0002 | 0.0365 |
| SPN | 3.95 | 1.55 | 4.14 | 0.0002 | 0.0365 |
| CSF1R | 2.35 | 7.45 | 4.14 | 0.0002 | 0.0365 |
| EBI3 | 3.73 | 0.38 | 4.12 | 0.0003 | 0.0365 |
| MPEG1 | 2.72 | 5.44 | 4.12 | 0.0003 | 0.0365 |
| ARHGAP30 | 2.41 | 4.09 | 4.12 | 0.0003 | 0.0365 |
| GBGT1 | 2.36 | 3.31 | 4.12 | 0.0003 | 0.0365 |
| LAG3 | 3.36 | 1.38 | 4.12 | 0.0003 | 0.0365 |
| GNA15 | 2.78 | 3.19 | 4.12 | 0.0003 | 0.0365 |
| HNMT | 1.62 | 4.82 | 4.11 | 0.0003 | 0.037 |
| RNASE6 | 3.28 | 3.75 | 4.11 | 0.0003 | 0.037 |
| CD53 | 2.2 | 5.04 | 4.1 | 0.0003 | 0.0371 |
| APBB1IP | 2.54 | 4.59 | 4.09 | 0.0003 | 0.0375 |
| LCK | 3.73 | 1.22 | 4.09 | 0.0003 | 0.0375 |
| CARD16 | 2.64 | 3.19 | 4.09 | 0.0003 | 0.0375 |
| CCR2 | 5.11 | -0.53 | 4.08 | 0.0003 | 0.0382 |
| HAVCR2 | 2.65 | 3.85 | 4.07 | 0.0003 | 0.0383 |
| PLEK | 2.86 | 4.54 | 4.07 | 0.0003 | 0.0383 |
| TRIM5 | 1.65 | 3.61 | 4.06 | 0.0003 | 0.0383 |
| IGHA1 | 7.69 | 2.42 | 4.06 | 0.0003 | 0.0383 |
| TSPAN14 | 1.4 | 5.86 | 4.06 | 0.0003 | 0.0384 |
| AC002316.1 | -3.19 | 2.04 | -4.05 | 0.0003 | 0.0385 |
| RHOF | 2.9 | 2.19 | 4.05 | 0.0003 | 0.0385 |
| CD84 | 3.24 | 5.49 | 4.05 | 0.0003 | 0.0385 |
| GPR137B | 2.1 | 4.71 | 4.04 | 0.0003 | 0.0385 |
| GLRA3 | -5.37 | 0.93 | -4.04 | 0.0003 | 0.0385 |
| CD37 | 3.11 | 3.04 | 4.03 | 0.0003 | 0.0385 |
| NOL4L | 2.03 | 4.37 | 4.03 | 0.0003 | 0.0385 |
| IRF1 | 1.98 | 4.99 | 4.03 | 0.0003 | 0.0385 |
| RENBP | 2.28 | 3.11 | 4.03 | 0.0003 | 0.0385 |
| SKAP1 | 3.16 | 0.52 | 4.03 | 0.0003 | 0.0385 |
| LAIR1 | 3.15 | 4.36 | 4.02 | 0.0003 | 0.0385 |
| PCDHB2 | -3.64 | 1.79 | -4.02 | 0.0003 | 0.0385 |
| LAMP3 | 3.09 | 1.39 | 4.01 | 0.0004 | 0.0392 |
| CD3E | 5.06 | 1.77 | 4.01 | 0.0004 | 0.0392 |
| MYOM2 | -2.96 | 2.54 | -4 | 0.0004 | 0.0392 |
| RBM47 | 2.8 | 2.6 | 4 | 0.0004 | 0.0392 |
| SLAMF7 | 4.01 | 2.79 | 3.99 | 0.0004 | 0.0403 |
| PLXND1 | 1.23 | 7.87 | 3.99 | 0.0004 | 0.0403 |
| ABI3 | 2.71 | 3.4 | 3.98 | 0.0004 | 0.0403 |
| PPARG | 2.57 | 3.95 | 3.98 | 0.0004 | 0.0403 |
| SPI1 | 2.89 | 5.41 | 3.97 | 0.0004 | 0.0409 |
| SLC29A4 | 3.04 | 2.64 | 3.97 | 0.0004 | 0.0414 |
| SNX20 | 4.9 | 0.65 | 3.96 | 0.0004 | 0.0417 |

| | | | | | |
|---|---|---|---|---|---|
| FRMD4A | 1.24 | 5.44 | 3.95 | 0.0004 | 0.0424 |
| IGSF6 | 2.92 | 3.02 | 3.95 | 0.0004 | 0.0425 |
| ZAP70 | 4.24 | 0.81 | 3.94 | 0.0004 | 0.0425 |
| CTSH | 2.03 | 5.68 | 3.94 | 0.0004 | 0.0425 |
| SETBP1 | -3.2 | 4.97 | -3.93 | 0.0004 | 0.0437 |
| CD8A | 4.78 | 1.1 | 3.93 | 0.0004 | 0.0437 |
| OSBPL1A | -2.09 | 4.13 | -3.92 | 0.0004 | 0.044 |
| GTF3C4 | -1.37 | 4.89 | -3.92 | 0.0005 | 0.0448 |
| HLA-DMB | 3.52 | 3.87 | 3.91 | 0.0005 | 0.0451 |
| HLA-DPA1 | 2.63 | 7.26 | 3.91 | 0.0005 | 0.0454 |
| CARMIL1 | -2.51 | 3.89 | -3.9 | 0.0005 | 0.0457 |
| HOXB5 | 2.96 | 0.4 | 3.9 | 0.0005 | 0.046 |
| RPP30 | 0.94 | 4.02 | 3.89 | 0.0005 | 0.0462 |
| UBR7 | -1.12 | 5.74 | -3.89 | 0.0005 | 0.0464 |
| LY9 | 3.32 | -0.33 | 3.88 | 0.0005 | 0.0464 |
| CACNA1B | -6.12 | -0.28 | -3.88 | 0.0005 | 0.0464 |
| TESPA1 | 3.4 | -0.11 | 3.87 | 0.0005 | 0.0481 |
| BARX1 | -6.7 | 1.64 | -3.86 | 0.0005 | 0.0492 |
| SORCS2 | -4.17 | 1.32 | -3.85 | 0.0005 | 0.0494 |
| RNF144B | -1.64 | 3.76 | -3.85 | 0.0006 | 0.0494 |
| IL2RA | 4.42 | 0.28 | 3.85 | 0.0006 | 0.0494 |

Supplemental Table 3

| Cell Type | Positive Markers | Negative Markers |
|---|---|---|
| osteoblast | ALPL, SP7, COL1A1, SPP1, SPARC, IBSP, RUNX2 | TOP2A, MKI67, SOX9, ACAN, SOST, MEPE, OSCAR, CD3D, VWF, CD14 |
| osteoblast_immature | ALPL, RUNX2 | SPP1, SPARC |
| osteoblast_proliferating | TOP2A, MKI67, RUNX2, ALPL, SP7 | |
| osteocyte | SOST, DMP1, FGF23, MEPE, PDPN, PHEX | ALPL, SP7, SPP1 |
| osteoclast | MMP9, CTSK, ACP5, CTR, OSCAR | SP7, ALPL |
| chondrocyte | ACAN, COL2A1, SOX9, COL9A1 | SP7, ALPL |
| msc | ENG, NT5E, THY1 | SP7, ALPL, CD34, PTPRC, CD19, CD14 |
| myeloid | CD74, CD14, LYZ, CD68 | CD34, CD3G, CD7, CD19, NCAM1, SP7, ALPL |
| TILS/NK | CD2, CD3D, CD3E, CD3G, GNLY, NKG7, KLRB1 | CD19, CD34, CD14, SP7, ALPL |
| endothelial | PECAM1, VWF, CDH5 | SP7, ALPL |
| naïve B-cells | MS4A1, CD19, PAX5, IGHD, IGHM | CD38 |
| pDC | LILRA4, CLEC4C, NRP1 | |
| plasmacells | CD27, SDC1, SLAMF7, CD74, CD38, IGHM | |

Supplemental Table 4

| Primary Antibodies | Product Details | Company/Ref Code | Primary Antibody Dilution | Secondary Antibody Dilution |
|---|---|---|---|---|
| pSTAT1 | p-Stat1 (Tyr701) (58D6) Rabbit mAb | CST #9167 | 750 | 5000 |
| STAT1 (human) | Stat1 (9H2) Mouse mAb | CST #9176 | 750 | 5000 |
| pTBK1 | p-TBK1/NAK (Ser172) (D52C2) XP® Rabbit mAb | CST #5483 | 750 | 5000 |
| TBK1 | TBK1/NAK (D1B4) Rabbit mAb | CST #3504 | 750 | 5000 |
| pIRF3 (human) | p-IRF3 (S386) (EPR2346) Rabbit mAb | Abcam #76493 | 1000 | 5000 |
| IRF3 | IRF-3 (D6I4C) XP® Rabbit mAb | CST #11904 | 1300 | 5000 |
| cGAS | cGAS (D1D3G) Rabbit mAb | CST #15102 | 1000 WB/200 IF | 5000 |
| pSTING (human) | p-STING (Ser366) (D7C3S) Rabbit mAb | CST #19781 | 1000 | 5000 |
| pSTING (mouse) | p-STING (Ser365) (D8F4W) Rabbit mAb | CST #72971 | 1000 | 5000 |
| STING | STING (D2P2F) Rabbit mAb | CST #13647 | 750 | 5000 |
| Tubulin | α/β-Tubulin Rabbit mAb | CST #2148 | 2500 | 10000 |
| Actin (human) | Actin Mouse mAb | CST #3700 | 2500 | 10000 |
| Actin (mouse) | Actin Rabbit mAb | CST #4967 | 2500 | 10000 |

Supplemental Table 5

| Target | Antibody | Company | Ref code | Dilution |
|---|---|---|---|---|
| CD45 | PerCP/Cy5.5 Rat Anti-Mouse CD45 [Clone: 30-F11] | Biolegend | 103132 | 1/200 |
| CD90.2 | BV785 Rat anti-Mouse CD90.2 [Clone: 30-H12] | Biolegend | 105331 | 1/200 |
| PD1 | PE Rat Anti-Mouse CD279 (PD-1) [Clone: 29F.1A12] | Biolegend | 135205 | 1/200 |
| CD4 | BUV395 Rat Anti-Mouse CD4 [Clone: GK1.5], BD Horizon | BD Biosciences | 565974 | 1/200 |
| CD8a | BUV737 Rat Anti-Mouse CD8a [Clone: 53.6.7], BD Horizon | BD Biosciences | 612759 | 1/200 |
| CD25 | APC/Cy7 Rat Anti-Mouse CD25 [Clone: PC61] | BD Biosciences | 552880 | 1/100 |
| Ly108 | APC Anti-Mouse Ly108 [Clone: 330-AJ] | Biolegend | 134609 | 1/200 |
| CD69 | PE-Cy7 Armenian Hamster Anti-Mouse CD69 [Clone: H1.2F3] | BD Biosciences | 552879 | 1/200 |
| CD103 | BV421 Armenian Hamster Anti-Mouse CD103 [Clone: 2E7] | Biolegend | 121421 | 1/200 |
| CD11b | BV605 Rat Anti-Mouse CD11B [Clone: M1/70] | BD Biosciences | 563015 | 1/200 |
| CD11c | BV650 Armenian Hamster Anti-Mouse CD11c [Clone: N418] | Biolegend | 117339 | 1/100 |
| Ly6c | BV711 Rat Anti-Mouse Ly-6C [Clone: HK1.4] | Biolegend | 128037 | 1/200 |
| CD45R | BV785 Rat Anti-Mouse/Human CD45R/B220 [Clone: RA3-6B2] | Biolegend | 103246 | 1/100 |
| Ly6G | BV785 Rat Anti-Mouse Ly-6G [Clone: 1A8] | Biolegend | 127645 | 1/200 |
| NK1.1 | BV785 Mouse Anti-Mouse NK-1.1 [Clone: PK136] | Biolegend | 108749 | 1/200 |
| SiglecF (CD170) | BV786 Rat Anti-Mouse Siglec-F [Clone: E50-2440], BD Optibuild | BD Biosciences | 740956 | 1/200 |
| F4/80 | AF647 Rat Anti-Mouse F4/80 [Clone: BM8] | Biolegend | 123122 | 1/200 |
| IAIE | Alexa Fluor® 700 Anti-Mouse I-A/I-E [M5/114.15.2] | Biolegend | 107622 | 1/100 |
| CD24 | PE-Cy7 Rat Anti-Mouse CD24 [Clone: M1/69] | Biolegend | 101822 | 1/200 |
| CD206 (MMR) | FITC Rat Anti-Mouse CD206 (MMR) [Clone: C068C2] | Biolegend | 141704 | 1/200 |
| CD49b | PE Armenian Hamster Anti-Mouse CD49b [Clone: HMα2] | Biolegend | 103506 | 1/200 |
